# Polygenic Risk for Skin Autoimmunity Impacts Immune Checkpoint Blockade in Bladder Cancer

**DOI:** 10.1101/2019.12.18.881805

**Authors:** Zia Khan, Flavia Di Nucci, Antonia Kwan, Christian Hammer, Sanjeev Mariathasan, Vincent Rouilly, Jonathan Carroll, Magnus Fontes, Sergio Ley Acosta, Ellie Guardino, Haiyin Chen-Harris, Tushar Bhangale, Ira Mellman, Jonathan Rosenberg, Thomas Powles, Julie Hunkapiller, G. Scott Chandler, Matthew L. Albert

## Abstract

PD-1 and PD-L1 act to restrict T-cell responses in cancer and contribute to self-tolerance. Consistent with this role, PD-1 checkpoint inhibitors have been associated with immune-related adverse events (irAEs), immune toxicities thought to be autoimmune in origin. Analyses of dermatological irAEs have identified an association with improved overall survival (OS) following anti-PD-(L)1 therapy, but the factors that contribute to this relationship are poorly understood. We collected germline whole genome sequencing data from IMvigor211, a recent phase 3 randomized controlled trial comparing atezolizumab (anti-PD-L1) monotherapy to chemotherapy in bladder cancer. We found that high vitiligo, high psoriasis, and low atopic dermatitis polygenic risk scores (PRSs) were associated with longer OS under anti-PD-L1 monotherapy as compared to chemotherapy, reflecting the Th17 polarization of these diseases. PRSs were not correlated with tumor mutation burden, PD-L1 immunohistochemistry, nor T-effector gene signatures. Shared genetic factors impact risk for dermatological autoimmunity and anti-PD-L1 monotherapy in bladder cancer.

## Introduction

PD-1 checkpoint inhibitors have made significant advances in metastatic urothelial carcinoma (mUC). Measures of pre-existing immunity have been associated with response and survival during anti-PD-(L)1 treatment (1). These measures capture tumor foreignness (i.e., neo-epitopes), the accessibility of the tumor to cytotoxic T-cells, and tumor or immune cell expression of PD-L1. These measures, however, only capture local tumor immunity. In mice, systemic immunity is required for successful tumor rejection by PD-1 checkpoint blockade and substantial evidence supports the role for host factors in influencing systemic immunity (2, 3). These host factors may include genetic variants that influence innate and adaptive immune responses, including risk for autoimmunity.

The PD-1 immune checkpoint acts to restrict T-cell responses and contributes to the regulation of immune self-tolerance (4, 5). Consistent with this role, patients develop immune related adverse events (irAEs) that are thought to be autoimmune in origin, but the link is poorly understood (6, 7). Indeed, in mice, on vulnerable genetic backgrounds spontaneous autoimmunity develops on knockout of PD-1. Yet, whether genetic background is relevant to development of irAEs in humans has not been tested. irAEs also vary in their frequency and organ system affected. Among the most common irAEs occur in the skin. Intriguingly, dermatological irAEs have also been associated with longer overall survival across PD-1 checkpoint inhibitors (8–12). To gain further insight into this relationship, we used genetic variants known to impact risk of psoriasis, vitiligo, and atopic dermatitis ascertained in independent cohorts by genome wide association study (GWAS). Using these variants, we computed polygenic risk scores for patients treated with atezolizumab or chemotherapy in IMvigor211 and associated them with dermatological irAE occurrence and overall survival.

## Results

To examine the relationship between safety and efficacy of checkpoint blockade, we conducted an analysis of irAEs using data from safety evaluable patients (N=459) receiving atezolizumab in IMvigor211(13). We grouped irAEs using an organ and system-based classification (see **Methods**). Skin irAEs were the most common, followed by gastrointestinal (GI) and endocrine irAEs (**Fig. 1a**). We confirmed a similar distribution of irAE in IMvigor210 (N=429), a phase 2 single-arm trial for treatment of mUC(14, 15). We focused on low grade irAEs as they are typically managed without systemic corticosteroids, and rarely lead to removal of patients from treatment(16). To address survival bias, we used a time-dependent covariate in a Cox proportional hazards model, including baseline factors as additional covariates (see **Methods**). In agreement with previous reports linking dermatological irAEs to survival(8–12), we found that overall survival was associated with low grade skin irAEs in IMvigor211 (p = 0.024; HR 0.66; 95% CI 0.45-0.95) and IMvigor210 (p = 0.0023; HR 0.53; 95% CI 0.35-0.80; **Fig. 1b**). To verify the robustness of our results, we conducted a landmark analysis. We selected landmarks at the point at which 90% patients in the low grade skin irAE group had experienced their event (**Fig. 1c**) and confirmed an association with improved OS in both trials (**Fig. 1d**).

**Figure 1.**
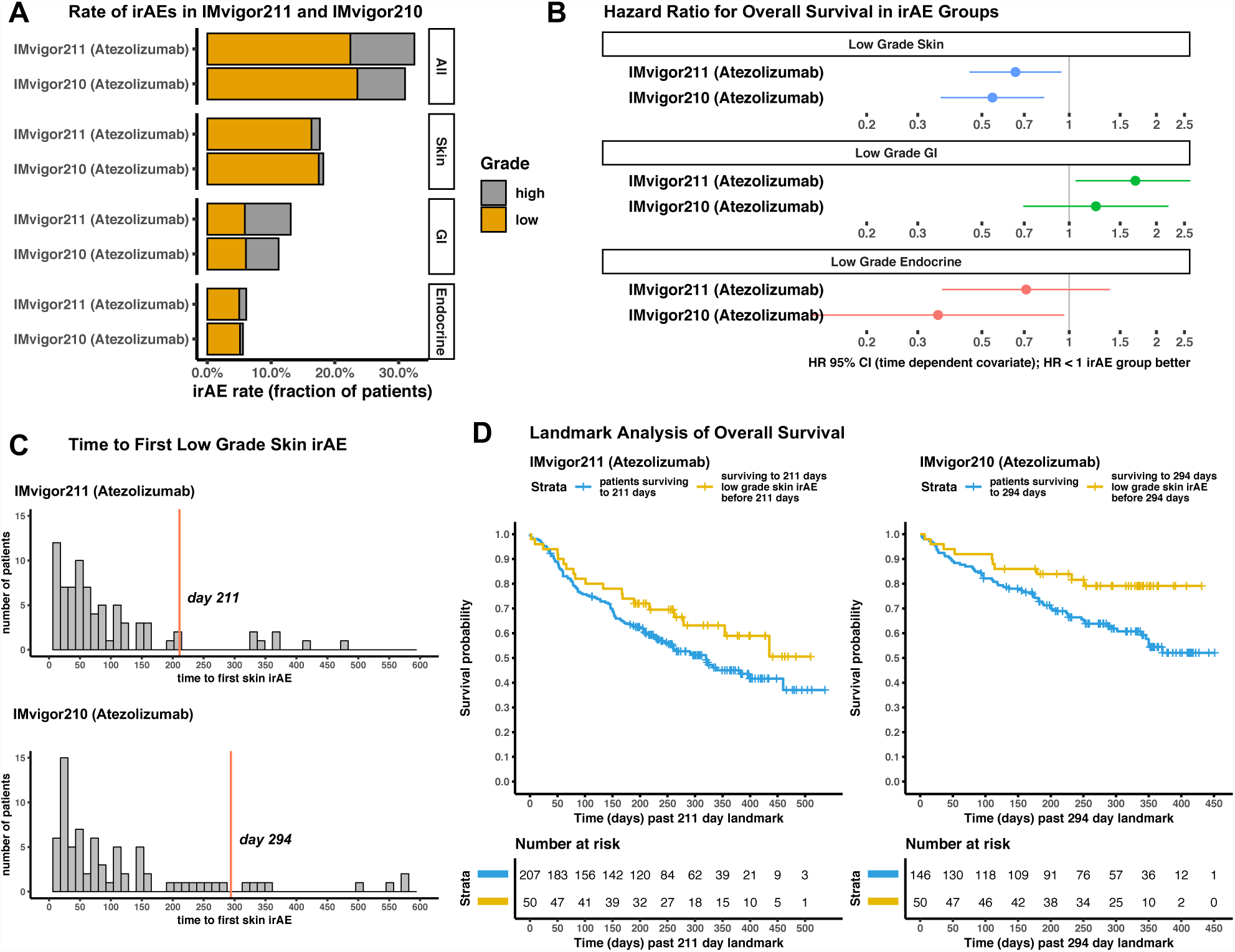
Low grade dermatological immune related adverse events (irAEs) in patients receiving atezolizumab are associated with longer overall survival. (a) Rate of irAEs, aggregated by system and organ-based classification, across trials. Only irAE categories with occurrence rates >5% are shown. Low = grade 1 or 2 (orange stacking bar); High = grade 3, 4, or 5 (gray stacking bar). (b) Hazard ratios (HRs) with 95% confidence intervals (CIs) are shown as thick lines, comparing overall survival of individuals that experienced low grade irAEs to those that did not experience irAE of a given classification. Individuals that experienced a high grade irAE of the given classification were excluded from the analysis. A time dependent covariate in a Cox proportional hazards model was used to estimate the HRs (see Methods). (c) Distribution of time to first skin irAE in days across trials. 90% of patients that experienced skin irAEs lie to the left of the orange landmark line. (d) Kaplan-Meier survival curves comparing the overall survival (OS) of individuals in the atezolizumab arm of IMvigor211 and IMvigor210 after a defined landmark. Tick marks show censoring events. GI = gastrointestinal.

30× germline whole genome sequencing data from 465 individuals within the IMvigor211 study (238 receiving atezolizumab and 227 chemotherapy treated) met strict filters for population and genotype data quality control (see **Methods**). We confirmed that OS and response rates were no different in these individuals, as compared to the intent-to-treat (ITT) population of 931 individuals (**Fig. S1**). We also confirmed that immune (IC) and tumor cell (TC) staining of PD-L1 by immunohistochemistry (IHC) and tumor mutation burden (TMB) were similarly balanced across trial (**Fig. S2**).

To test the hypothesis that genetic factors shared with dermatological autoimmunity impact anti-PD-L1 safety and efficacy, we used publicly available GWAS summary statistics to construct PRSs for psoriasis (PSO), atopic dermatitis (AD), and vitiligo (VIT) which were ascertained on independent case/control cohorts (see **Methods, Fig. 2a, Table S1, Fig. S3**) (17, 18). We used two studies for psoriasis, Immunochip (PSO/IC) and the UK Biobank (PSO/UKBB). PSO/IC assessed genotypes curated for immunologically relevant variants, whereas psoriasis cases in PSO/UKBB were self-reported and assayed using genome-wide genotyping. We additionally constructed PRSs for Alzheimer’s disease (ALZ) to serve as negative controls. Controlling for genotype principal components (PCs) and gender, we identified associations at a false discovery rate (FDR) of 10% between the occurrence of skin irAEs and genetic risk for psoriasis within the atezolizumab arm of IMvigor211 (**Fig. 2b, Table S2**). The associations were consistent across GWASs used to construct the psoriasis PRSs (**Fig. 2c**).

**Figure 2.**
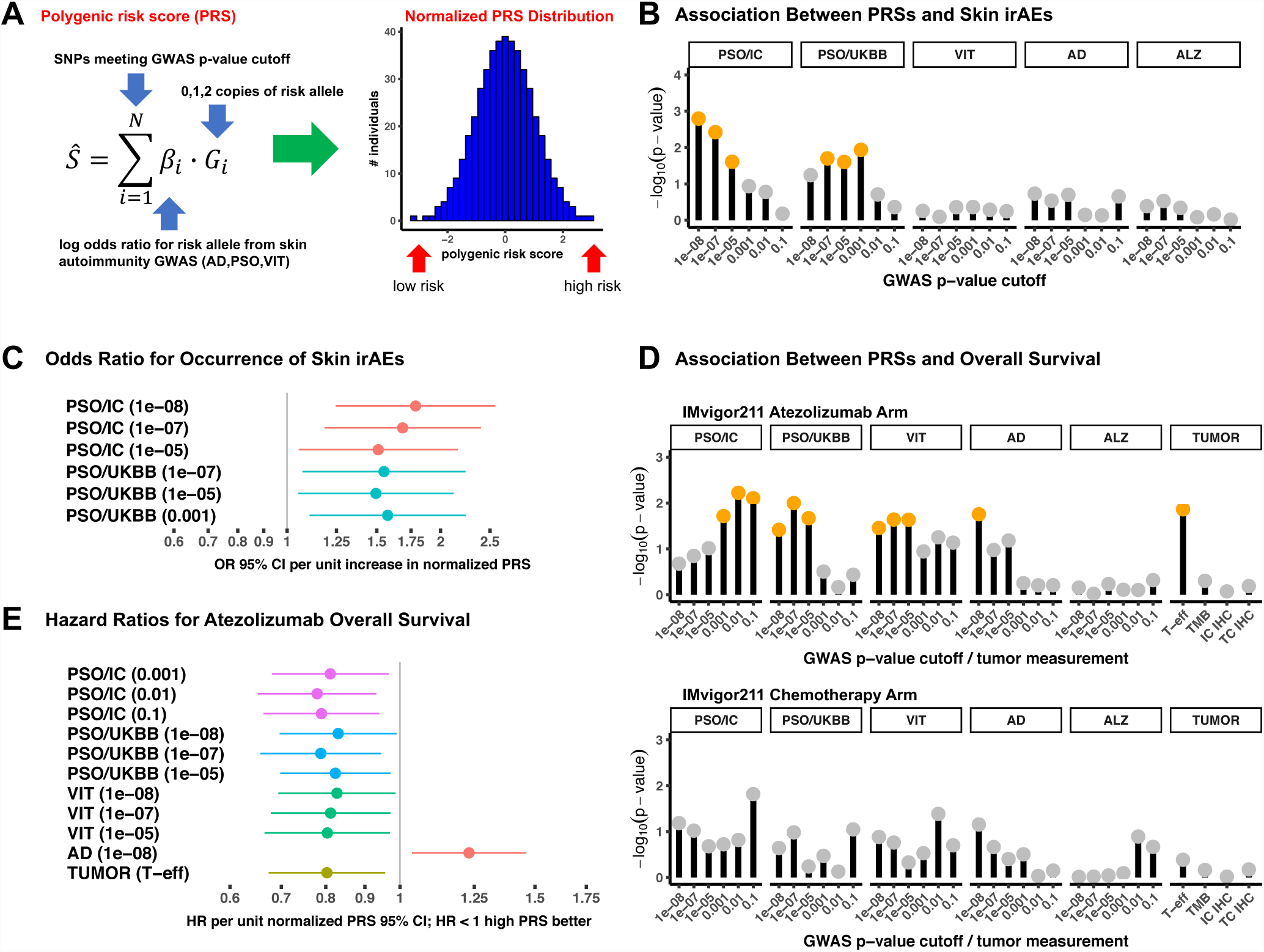
Polygenic risk for skin autoimmunity is associated with the occurrence of dermatological irAEs and overall survival in the atezolizumab arm of IMvigor211. (a) Polygenic risk scores (PRSs) were computed for each individual with whole genome germline sequencing data in IMvigor211 for dermatological autoimmune diseases: atopic dermatitis (AD), psoriasis (PSO), and vitiligo (VIT). PRSs were constructed for 6 p-value cutoffs (see Table S2 and Methods). (b) Negative log_10_ p-values for a given PRS testing for association with occurrence of skin irAEs controlling for 5 genotype principal components and gender by logistic regression. Orange circles show associations that were significant at false discovery rate (FDR) of 10%. Gray circles did not meet statistical significance. (c) Odds ratios (ORs) and 95% confidence intervals (CIs) were estimated for PRSs and skin irAE occurrence for PRSs that met a significance cutoff of FDR 10%. ORs are expressed in per unit change of a normalized PRS. GWAS p-value cutoff indicated within parentheses. (d) Negative log_10_ p-values for a given GWAS and p-value cutoff PRS testing for association with overall survival using a Cox proportional hazards model controlling for 5 genotype principal components and baseline clinical factors (see Methods). (e) Adjusted hazards ratios (HRs) and 95% CIs for PRS and OS associations are shown, using significance cutoff of 10% FDR. HRs are expressed in unit change of a normalized PRS. IC = Immunochip; UKBB = UK Biobank; T-eff = CD8 T-effector gene expression signature score; TMB = tumor mutation burden; IHC = immunohistochemistry; IC IHC = PD-L1 expression on immune cells; TC IHC = PD-L1 expression on tumor cells.

Given the observations that dermatological irAEs are associated with longer OS, and germline genetic risk for psoriasis is associated with increased odds of skin irAEs during checkpoint blockade, we investigated whether dermatological autoimmune disease PRSs were associated with OS under atezolizumab treatment, or chemotherapy (analyzed independently). PRSs for atopic dermatitis, psoriasis, and vitiligo were associated with OS in the atezolizumab arm, but not the chemotherapy arm, of IMvigor211 at an FDR of 10%, controlling for baseline clinical factors in addition to genotype PCs (**Fig. 2d, Table S2**; see **Methods**). PRSs for Alzheimer’s disease, as expected, did not show any significant association. The GWAS p-value cutoffs at which we observed significant associations differed between OS and skin irAEs, as well as across diseases, reflecting the differing degrees of genetic sharing and statistical power of the GWASs underlying the PRSs (see **Supplemental Discussion**). We also highlight that the T-effector gene signature, a measure of CD8+ T-cell effector function within a tumor, was the unique tumor factor significantly associated with OS within the atezolizumab arm in the IMvigor211 (confirming prior studies (14)).

Both increased polygenic risk for vitiligo and psoriasis were associated with longer OS under treatment with atezolizumab, whereas decreased polygenic risk for atopic dermatitis was associated with longer OS under atezolizumab treatment (**Fig. 2e**). We confirmed that the OS associations were not simply due to correlation between the T-effector signature or strong intercorrelation among the disease-specific PRSs (**Fig. S4**). We quantile normalized the T-effector signature score to allow comparison to quantile normalized PRSs. The hazard ratio benefit of a higher T-effector signature score per normalized unit was 0.81, similar to that of the PRSs per normalized unit for psoriasis, vitiligo, and the inverse of atopic dermatitis 0.78-0.83.

We then assessed whether PRSs were prognostic, informative of outcome regardless of treatment, or predictive, informative of the effect of experimental treatment. To formally test if PRSs were predictive, we incorporated both trial arms into a Cox proportional hazards model, and assessed interaction between the PRS and trial arm. After controlling for baseline clinical covariates and genotype PCs, a significant trial arm by PRS interaction term tests whether the hazard ratio comparing atezolizumab to chemotherapy differs between patients with high versus low PRSs (see **Methods**). At an FDR of 10%, we found that PRSs for atopic dermatitis, psoriasis and vitiligo were predictive of OS in IMvigor211 (**Fig. 3a, Table S2**). Consistent with prior reporting of the IMvigor211 study, tumor measurements were not strongly predictive of OS (13).

**Figure 3.**
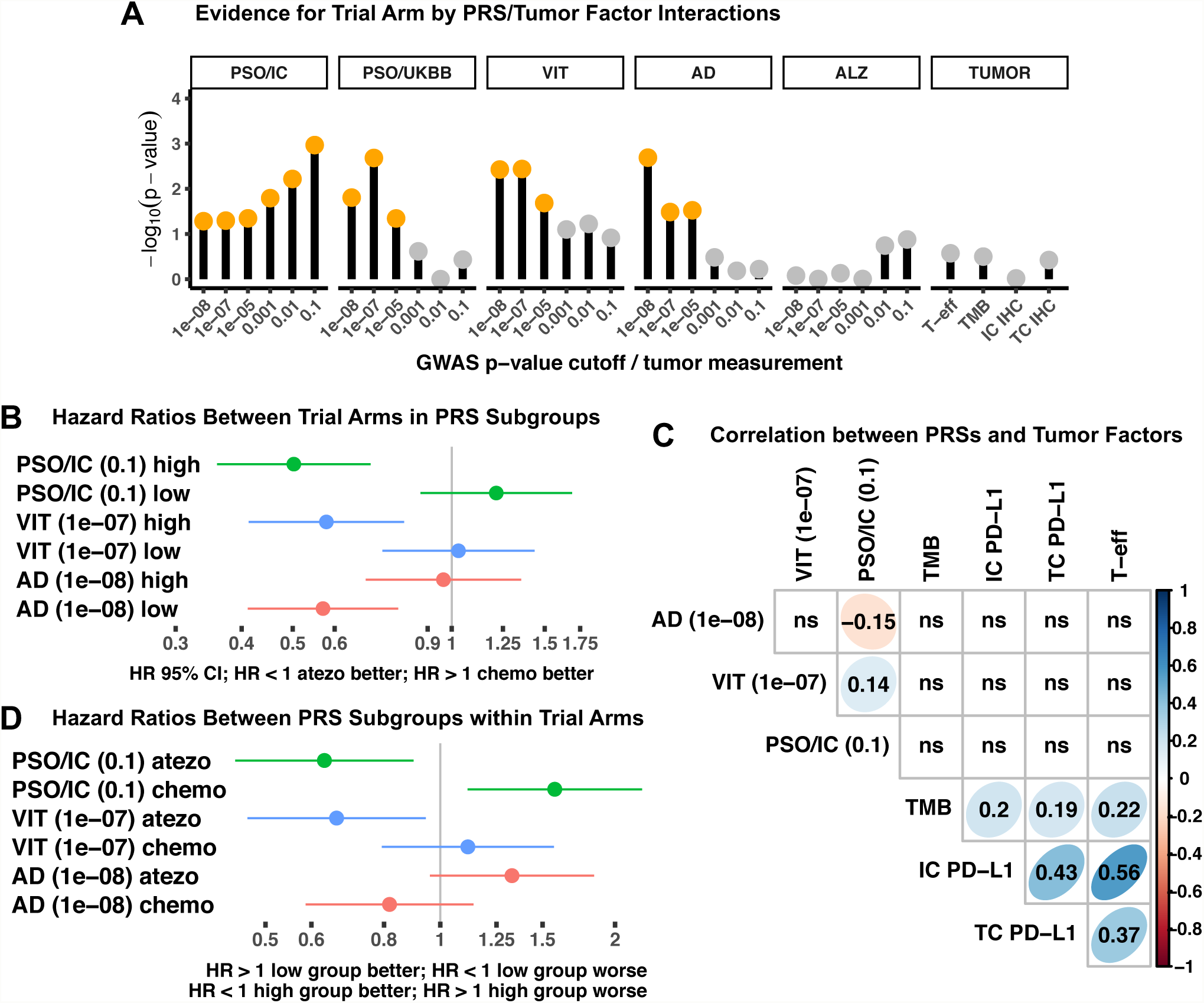
Polygenic risk scores for dermatological autoimmunity are informative of the effect of treatment in IMvigor211. (a) Negative log_10_ p-values for a given PRS testing for a statistically significant trial arm by PRS/tumor factor interaction using a Cox proportional hazards model for overall survival, controlling also for 5 genotype PCs, gender and clinical covariates (see Methods). Orange circles designate PRS and trial arm interaction terms that were significant at an FDR of 10%. Gray circles indicate values that did not meet statistical significance. (b) HRs and 95% CI for Cox proportional hazard models are shown, comparing trial arms in subgroups of high and low risk for PRSs that showed the strongest trial arm by PRSs interaction. HRs were adjusted for baseline clinical covariates, genotype PCs and gender (see Methods). High and low risk groups were defined by splitting the population on the median. Respective GWAS p-value cutoff of the corresponding PRS are indicated in parentheses. (c) Spearman’s rank correlation between PRSs and tumor factors across individuals in IMvigor211. Rank correlations with p ≥ 0.05 are labeled ns. T-eff = CD8 T-effector gene expression signature score; TMB = tumor mutation burden; IC PD-L1 = immune cell PD-L1 expression by IHC; TC PD-L1 = tumor cell PD-L1 expression by IHC. (d) HRs and 95% CIs from Cox proportional hazard models are shown comparing OS of subgroups of high and low risk for PRSs within each trial arm. atezo = atezolizumab; chemo = chemotherapy.

To better understand the behavior of the PRSs, we split the population on median PRS, creating two subgroups of individuals with high and low polygenic risk. We focused on the respective PRSs that had the strongest trial arm by risk score interaction for each dermatological autoimmune disease. High vitiligo (p = 0.0016; HR 0.58; 95% CI 0.41-0.81) and high psoriasis risk (p = 5.5×10-5; HR 0.50; 95% CI 0.36-0.70) individuals had better OS under checkpoint blockade than chemotherapy, whereas low atopic dermatitis risk (p = 0.0008; HR 0.57; 95% CI 0.41-0.79) individuals had improved OS under atezolizumab treatment as compared to chemotherapy (**Fig. 3b; Fig. S5-S7**). PRSs were uncorrelated tumor immune cell staining of PD-L1 by IHC, tumor mutation burden, and across patients (**Fig. 3c**). To gain further insight into the PRSs, we compared high and low PRS subgroups within each treatment arm (**Fig. 3d**). High vitiligo risk reflected improved OS in the atezolizumab arm, whereas high psoriasis risk and low AD risk groups reflected both improved OS in the atezolizumab arm and worse OS within the chemotherapy arm. As response is a proxy for longer and shorter OS, we found that the response rates within these subgroups followed a similar numeric pattern (**Fig. S8**).

Variants in the MHC locus and specific HLA alleles contribute significantly to genetic risk for psoriasis, vitiligo, and atopic dermatitis (**Table S3**) (19–21). As our PRSs were generated using a reference panel that poorly approximated linkage disequilibrium in the MHC region, we excluded variants from this region in our association analyses. To address this caveat, we repeated our analysis including variants in the MHC region. Inclusion of this additional information did not strengthen the associations we observed (**Fig. S9**). We also called HLA alleles on the basis of direct sequence evidence (see **Methods**). We found that HLA alleles previously found to be associated with risk of psoriasis, atopic dermatitis, or vitiligo were not associated with OS in the atezolizumab or the chemotherapy arm of IMvigor211 (**Table S4**). As these alleles may contribute additively to risk of dermatological autoimmunity, we incorporated the risk conferred by these alleles to our PRSs (see **Methods**). Inclusion of this additional information had negligible impact (**Fig. S9**).

The tumor microenvironment (TME) consists of a complex mixture of immune cells, stromal, and tumor cells (22). Polygenic risk may influence the composition of the TME, which in turn, impacts patient survival. Using pre-treatment, bulk tumor gene expression for N=398 individuals with both RNA-seq and germline genetic data, we generated immune and stromal cell type enrichment scores and, to limit the multiple testing burden, we associated them with PRSs that had the strongest trial arm by risk score interaction for each dermatological autoimmune disease(23). We found no evidence for association between cell type enrichment scores and PRSs (**Fig. S10**). Alternatively, PRSs might be relevant only in certain tumor contexts. To address this question, we delineated four subgroups on the basis of high or low PRS and a high or low tumor factor. Specifically, we considered: PD-L1 expression measured by IHC and expressed on immune or tumor cells; TMB; and the CD8 T-effector signature. We additionally considered the tumor expression of selected T-helper chemokines and cytokines involved in differentiation, recruitment, and response (**Table S5**). We statistically assessed whether combining a PRS and a tumor factor was informative of the treatment effect on overall survival than the PRS alone (see **Methods**). This would occur if the treatment effect depended on both the PRS value and the tumor factor value (**Fig. S11**). We found high psoriasis PRS was beneficial in immune infiltrated tumors as reflected by survival benefit in tumors with high values of immune cell (IC) staining of PD-L1 by IHC and high expression of genes involved in CD8+ T-effector function. Although little or no pre-treatment IL17A/F expression was detected by bulk RNA-seq, genes involved in Th17 function including *IL23A, CXCL2* and *CCL20* also delineated a subgroup that strongly benefited from atezolizumab as compared to chemotherapy (**Fig. 4**). This subgroup existed in the absence of any correlation between psoriasis PRS and these tumor factors (**Fig. S12**). We also observed a divergent pattern between psoriasis PRS and expression of subunits of IL-12 (*IL12A* and *IL12B*) in a manner consistent with preclinical observations of the divergent roles of IL-12 and IL-23 in autoimmunity (see **Supplemental Discussion**; **Fig. S13-S14**).

**Figure 4.**
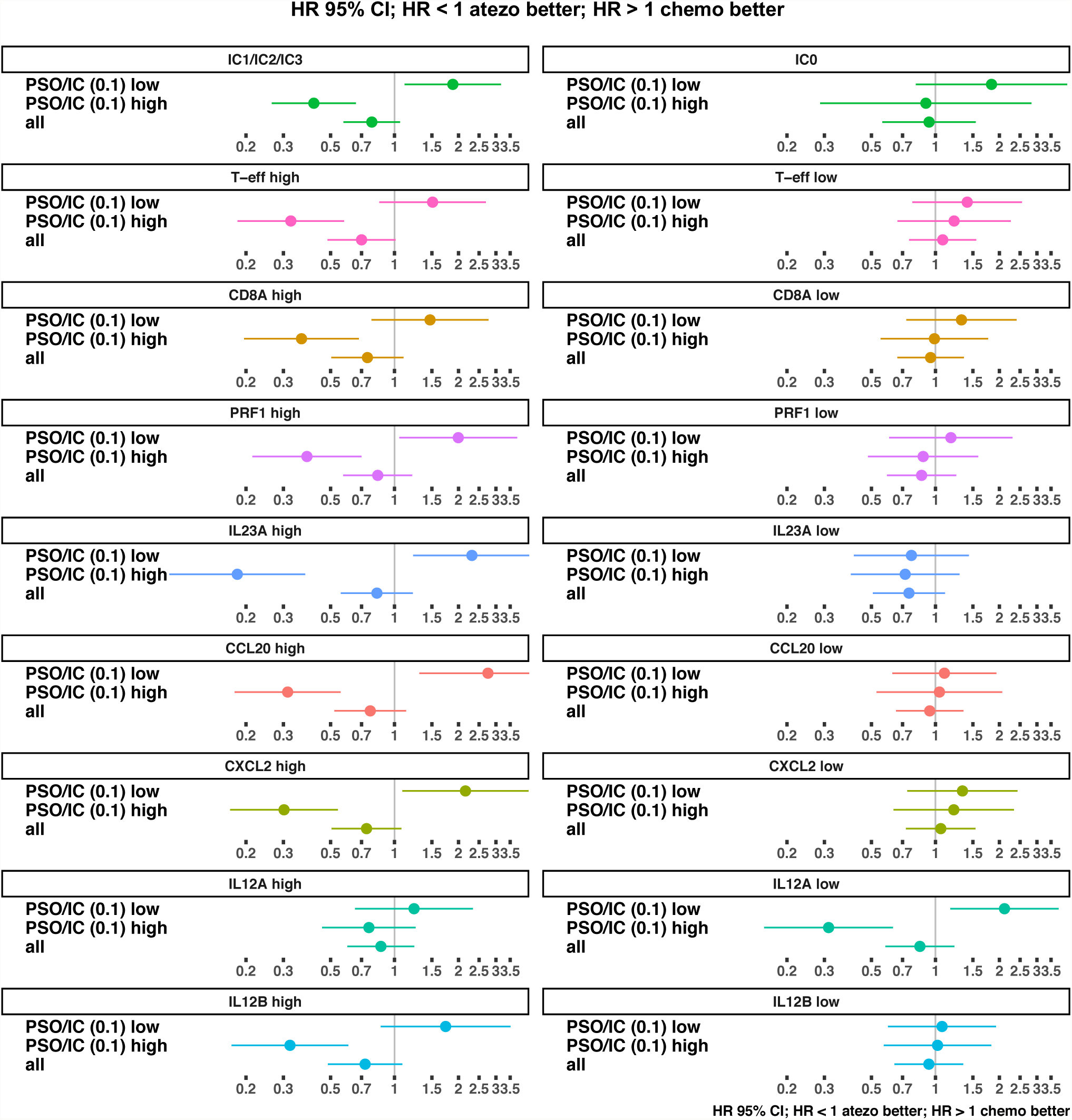
Polygenic risk for psoriasis is informative of the effect of treatment in specific tumor immune contexts. HR and 95% CI are shown, comparing trial arms in stratified by of high or low tumor factor, as shown, and stratified high or low risk for psoriasis using the PSO/IC PRS at a GWAS p-value cutoff of 0.1 where “all” designates no PRS stratification. HRs were adjusted for RNA-seq batch effects, baseline clinical covariates, genotype PCs, and gender (see Methods). High or low PRS groups and high or low tumor gene expression groups were defined by splitting the population on their median values, respectively. IC0-3 designate increasing levels of immune cell (IC) staining of PD-L1 by immunohistochemistry. Only tumor factors and genes that met defined filtering criteria (see Table S5) and were significant at an FDR of 10% are shown here (see Fig. S10 for all tested PRSs, genes, and tumor factors). atezo = atezolizumab; chemo = chemotherapy

## Discussion

Dermatological irAEs have been associated with longer overall survival across PD-1 checkpoint inhibitors (8–12). Using germline whole genome sequencing data from a phase 3 trial comparing anti-PD-L1 monotherapy to chemotherapy in mUC, we found that PRSs for dermatological autoimmune diseases were associated with increased risk for skin irAEs and longer OS with atezolizumab, as compared to chemotherapy. Our study provides further insight into the relationship between dermatological irAEs and patient survival. Our analysis indicates that genetic factors that modify individual risk of dermatological autoimmunity also impact PD-L1 blockade in this setting. Further study of the mechanisms and genes by which these variants act may provide important insights for the basis of novel cancer immunotherapies. The directionality of the association with OS and PRSs for psoriasis and vitiligo versus atopic dermatitis reflected the high and low Th17 polarization of these diseases in European populations, respectively (see **Supplemental Discussion**)(24, 25). Taken together, these observations support the notion that an immune set point, at least in part determined by germline genetic variation, may influence the efficacy of immune checkpoint blockade (1).

Although useful to provide insight into the role of autoimmunity during PD-L1 blockade, several important challenges remain before PRSs can contribute to clinical decision making. PRSs derived from GWASs conducted in European populations will have reduced accuracy in populations of Asian and African descent (26). PRSs are also limited by disease heritability and sample size of the underlying GWAS. Further studies are needed to establish the contexts where PRSs might be relevant and delineate their interactions with tumor factors in patients treated with immune checkpoint inhibitors.

## Materials and Methods

Detailed methods are described in the supplementary materials.

### Code Availability

An R package with anonymized clinical data, precomputed normalized PRSs, precomputed HLA calls for VIT, PSO, and AD risk alleles, and all R code to produce figures in the manuscript has been made available for download: http://research-pub.gene.com/CITSkinSurvival.

### Data Availability

De-identified patient level clinical data and summary level PRS data is available within the R package described above. All requests for anonymized genotype (VCF) or raw data (BAM/FASTQ) will be promptly reviewed by Roche/Genentech to verify if the request is subject to patient consent and confidentiality obligations. Any data that can be shared will be released through establishment of a data sharing agreement with Roche/Genentech. Please contact the corresponding authors for any inquiries regarding the requests for data.

## Acknowledgements

We thank all of our Genentech colleagues involved in the Human Genetics Initiative. We acknowledge Elaine Murray for help with curating irAE data sets and Felix Arellano for comments on early drafts of this manuscript. We acknowledge the Cancer Immunotherapy Committee (CITC), Genito-Urinary-Global Development Team (GU-GDT) for their support, the IMvigor210 and IMvigor211 study teams, the investigators, and the patients that contributed their samples and data for this study.

