## Supplemental Information for "Polygenic Risk for Skin Autoimmunity Impacts Immune Checkpoint Blockade in Bladder Cancer"

**This PDF file includes:**

Supplemental Discussion
Methods
Tables S1-S5
Figures S1-S14
Supplemental References

**Supplemental Discussion**

**Relationship Between Shared Genetics, GWAS statistical power, and PRS p-value cutoffs**

Significant associations between a PRS constructed from GWAS summary statistics and a phenotype in an independent cohort implied that shared genetic factors impact the disease risk studied in the original GWAS and the phenotype of interest(1). The GWAS p-value cutoff at which variants are considered for inclusion in the PRS impact the strength of the associations observed. We opted for a strategy whereby we tested several GWAS p-value cutoffs and reported significant associations at false discovery rate, thus accounting for multiple statistical tests (see **Methods**). For completeness, we report the results of each statistical test conducted. However, the following results pertaining to the GWAS p-value cutoff with the strongest association and the total number of significant associations requires further discussion:

- Only PSO/IC and PSO/UKBB PRSs are associated with the occurrence of skin irAEs (**Fig. 2b**).
- The p-value cutoff at which the PSO PRSs are most strongly associated skin irAEs differs from which the p-value cutoff at which PSO PRSs are most strongly associated with OS (**Fig. 2b, 2d**).
- The p-value cutoff that is most strongly associated with OS or predictive of OS differs across PRSs derived from different dermatological autoimmune diseases (**Fig. 2d, Fig. 3a**).
- The total number of GWAS p-value cutoffs at which significant associations are observed differs across dermatological autoimmune diseases and phenotypes (**Fig. 2b, 2d, Fig. 3a**).

There are three major factors that contribute to these results, none of which impact the statistical validity of our analysis and conclusions. First, the GWASs that underlie the PRSs vary in their statistical power. Statistical power is influenced by a number of factors, in addition to sample size of the GWAS, such as heritability and genetic architecture, definition of disease cases. This, in turn, will impact the p-value cutoff at which we observe a significant association. Some of these non-linear relationships can be explored in simulation using recently developed tools (https://github.com/andreyshabalin/simPRS). Second, we approximated linkage disequilibrium (LD) in the original GWASs using a reference panel to construct our PRSs (see **Methods**). How accurately this LD reference panel reflects LD in the original GWAS population will also impact the statistical power of the final PRS constructed. Third, the associations with PRSs imply shared genetics, but the extent of sharing is unknown. The extent of this sharing may differ between phenotypes and across diseases. The extent of this sharing might will also influence the strength of the associations and the GWAS p-value cutoff at which a significant PRS phenotype association is observed. If the sharing is broad, then stronger associations may be observed at higher, not lower, p-value cutoffs. If it is focused on a subset of GWAS disease risk variants, then we might expect significant associations to be observed at smaller GWAS p-value cutoffs.

Due to these considerations, significant associations at multiple p-value cutoffs do not necessary imply the results we observe are more robust than significant associations observed at a single, or fewer, p-value cutoffs. Associations observed despite differences in how GWAS cases were defined in the original studies does imply a degree of robustness. In the case of the two psoriasis PRSs, the definition of cases was achieved in a distinct manner: PSO/UKBB cases are self-reported; whereas PSO/IC was diagnosed by a physician. The observation of significant association with two polar Th17 diseases psoriasis and vitiligo and a negative association with atopic dermatitis does implies an additional degree of robustness. Significant PRS and phenotype associations were observed despite differing clinical definitions of these diseases.

**Relationship Between Th17 Cells and PD-(L)1**

PD-L1 and/or PD-1 have been directly linked to psoriasis pathogenesis in murine models. Skin responses of PD-1 knockout mice were more severe in an imiquimod induced model of psoriasis, as reflected by higher neutrophilic infiltration, Th17 cytokines and hyperplasia, as compared to wild-type controls (2). In experimental models, adoptive transfer of Th17 cells has been shown effective at rejecting tumors by activating tumor-specific $CD8^+$ T-cells (3). In humans, $CD4^+$ T-cells from anti-PD-1 responding patients were found to have higher production of IL17A on stimulation than individuals that were non-responders or not treated with anti-PD-1(4). Similarly, PD-1 blockade in conjunction with T-cell stimulation of peripheral blood from cancer patients has been shown to augment Th17 and suppress Th2 cytokine responses(5). Although Th17 cells and related cytokines can support tumor immunity, there exists evidence supporting a role for Th17 cells in promoting tumor growth(6). This may reflect the recruitment of neutrophils as a secondary response to inflammation(7). Following from our analyses, PD-(L)1 blockade might induce these mechanisms in individuals with high genetic polarization toward Th17 immunity, as reflected by high PRS for psoriasis or vitiligo.

**Divergent Pattern Between IL-12 Subunits**

In addition to pre-existing, CD8 T-effector function, we found that combining psoriasis PRS and the tumor expression level of genes involved in Th17 recruitment and response identified a subgroup that benefitted significantly from atezolizumab as compared to chemotherapy. While this supports the importance of Th17 driven immunity, it might arise out of intercorrelation among these tumor genes (see **Fig. S12**). Therefore, the divergent relationship between psoriasis PRS subunits of IL-12 (*IL12A* and *IL12B*) tumor expression was of particular interest (**Fig. 4**). The *IL12A* gene product IL12p35 combines with the *IL12B* gene product IL12p40 to form the IL12p70 heterodimer. IL12p40 also combines with the *IL23A* gene product IL23p19 to form IL-23 (8). Although IL12p70 has been associated with Th1 immunity, there is also evidence for its role in limiting late-stage autoimmune inflammation (9–11). Consistent with the latter mechanism of action, we found low *IL12A* to be associated with better outcome in patients possessing high psoriasis PRS. By contrast, high levels of both *IL23A* and *IL12B* were associated with better outcome in individuals with high psoriasis PRS (**Fig. 4; Fig. S13**). Additionally, *IL12A* showed a weaker correlation with $CD8^+$ T-effector signature genes, as compared to *IL12B* (**Fig. S14**). The directionality of the associations observed provides further support for the importance of Th17-driven effector mechanisms in anti-PD-L1 mediated tumor immunity.

**Methods**

**Patient Cohorts**

We conducted our analysis of immune related adverse events (irAEs) in the safety evaluable population from IMvigor211 and IMvigor 210 (12–14). The committees that approved study protocols and confirmation of informed consent from all study participants are included in the previous publications.

We sequenced DNA isolated from blood samples from N=479 individuals from IMvigor211 on the basis of availability and signed consent for research on germline DNA.

**Definition of irAE Categories**

irAE data conformed to an internal Adverse of Events of Special Interest (AESI) strategy. The strategy identified a collection of adverse event terms that had a putative immune-related etiology. We grouped AESIs into organ and system categories: skin, gastrointestinal (GI), and endocrine to allow meaningful statistical associations with survival. We grouped the following AESIs into the skin category: "Immune-Related Rash" and "Immune-Related Severe Cutaneous Reaction". The GI category consisted of the following AESIs: "Immune-Related Hepatitis," "Immune-Related Colitis," and "Immune-Related Pancreatitis." The endocrine category consisted of the following AESIs: "Immune-Related Hypothyroidism," "Immune-Related Hyperthyroidism," "Immune-Related Adrenal Insufficiency,", "Immune-Related Diabetes Mellitus," and "Immune-Related Hypophysitis." All other system and organ-based categories renal, neuro-muscular, and pulmonary, and systemic occurred at rates <5% and were not considered for association due to limited statistical power. Grading for AESIs was defined according to the National Cancer Institute Common Terminology Criteria for Adverse Events (CTCAE) as delineated in the original study protocols.

**Time-Dependent irAE Associations**

We used a time-dependent covariate in a Cox proportional hazards model that incorporated the time to irAE in our assessment of association of between survival and irAEs. Our analyses controlled for the following covariates, which were measured at baseline prior to treatment:

- Presence or absence of liver metastases.
- High or low CRP where (>10 mg/ml is high)
- Neutrophil to lymphocyte ratio normalized to the quantiles of the standard normal distribution.
- High or low alkaline phosphatase in serum at baseline (>147 IU/L was designated as high).
- High or low albumin in serum at baseline (<35g/L is low).
- High or low LDH in serum at baseline (>400 g/L is high).
- Gender
- Baseline ECOG status (0 or 1).
- Immune cell (IC) staining of PD-L1 by IHC. IHC data was obtained as using methods previously described in the original study publication and protocols (12–14).

Time dependent covariate was constructed using the tMerge function in the R survival package (https://cran.r-project.org/web/packages/survival/index.html). In contrast to landmark analysis, the use of a time-dependent covariate allowed us to control for the above additional covariates. All analyses, both the time-dependent covariate and the landmark approach, excluded individuals that experienced the high grade irAE from the non-irAE group for the irAE class that was analyzed.

**Whole Genome Sequencing**

Genomic DNA was extracted from blood samples using the DNA Blood400 kit (Chemagic) and eluted in 50µL Elution Buffer (EB, Qiagen). DNA was sheered (Covaris LE220) and sequencing libraries were prepared using the TruSeq Nano DNA HT kit (Illumina Inc.). Libraries were sequenced at Human Longevity (San Diego, CA, USA). 150bp paired-end whole-genome sequencing (WGS) data was generated to an average read depth of 30× using the HiSeq platform (Illumina X10, San Diego, CA, USA) and processed using the Burrows Wheeler Aligner (BWA)/Genome Analysis Toolkit (GATK) best practices pipeline (15–17). Short reads were mapped to hg38/GRCh38 (GCA_000001405.15), including alternate assemblies, using an alt-aware version of BWA to generate BAM files (18). All sequencing data was checked for concordance with SNP fingerprint data collected before sequencing.

**Genotype Data and Population Quality Control**

Only variants that passed GATK threshold for overall quality were used. Genotypes with the GATK-assigned quality (GQ) ≤ 20 were set to missing followed by removal of variants with an across sample missing genotype rate of 0.1 or more. Samples were also checked for a high rate of missing variant calls and extreme values of homozygosity as measured by inbreeding coefficient (F) estimates. None were removed by these filters. We also conducted pairwise identity-by-descent analysis (patient relatedness), no pair of samples were detected that had proportion IBD (PI_HAT > 0.1) that also had P(IBD=0)/Z0 of greater than 0.4 or more. We also asked if any one of the samples had a proportion IBD (PI_HAT > 0.1) with a large number (>200) of other samples indicating possible cross-contamination. No samples were filtered at this step. We performed PCA using EIGENSTRAT (19). Five rounds of PCA outlier removal iterations were performed at the default settings of EIGENSTRAT. Then, the final PCA was then performed to compute eigenvectors – top 5 of which were used in association analysis to correct for population stratification. The last filter removed a total of 14 samples.

**Construction of Polygenic Risk Scores**

All polygenic risk scores were constructed using publicly available GWAS summary statistics for vitiligo, psoriasis, and atopic dermatitis. We used a linkage disequilibrium (LD) reference panel, the European (EUR) population from the 1000 Genomes Project, to approximate LD that would be present in the original case control populations for each of the GWAS of these diseases. The whole genome sequencing data we collected was in GRCh38 coordinates with corresponding rsids from dbSNP v150. For each of the summary statistics, we remapped any rsids that changed from old versions of dbSNP to dbSNP v150. We used variants that were called within our IMvigor211 WGS data. We only used variants that had estimated odds ratios in the summary statistics. A few of the variants appeared as duplicates in the summary statistics, and in such instances, only the entry with the smallest p-value was kept. Variants with strand ambiguity (A/T or C/G genotypes) were removed. Variants that were not present in the EUR population in 1000 Genomes Project were also filtered out. We confirmed that the risk allele in the summary stats matched the alleles called at variants in our WGS data. Because we used WGS data, a small number of SNPs were multi-allelic in our WGS data. All other alleles were ignored and only the presence of the risk allele contributed to the PRS for the individual. LD clumping was performed using our LD reference panel to extract independent signals from the GWAS summary statistics (20). Specifically, a 250kb window around the index SNP was employed, adding variants with $r^2 > 0.25$ to the current clump (corresponding to PLINK parameters --clump-p1 1 --clump-p2 1 --clump-r2 0.25 --clump-kb 250). The PRS was computed as follows:

$$\hat{S} = \sum_{i=1}^{M} \beta_i \cdot G_i$$

where $M$ is the number of SNPs at a given GWAS p-value cutoff and $\beta_i$ corresponds to the log odds ratio of the $i$th SNP and $G_i = \{0,1,2\}$ corresponding to the number of copies of the risk allele. PRSs were quantile normalized to the quantiles of a standard normal distribution to allow comparison across GWAS.

Our risk scores did not include non-autosomal SNPs. Removal of non-autosomal SNPs assured that the total number of risk variants carried by any one individual would not differ on the basis of gender. Thus, scores from females would were on a comparable scale to scores generated from males. Our risk scores also did not include SNPs within the major histocompatibility complex (MHC) region nor did we include SNPs other difficult-to-genotype regions that have alternate contigs in the reference genome (alt regions in https://www.ncbi.nlm.nih.gov/grc/human). We excluded the MHC region due to the complex and strong LD patterns in this region which render it challenging to identify independent signals by LD clumping and because these complex LD patterns were poorly approximated by our LD reference panel. We explored the impact of excluding this region on our PRS associations in Figure S9. We also accounted for risk conferred by variants in this region by calling HLA alleles directly called from whole genome sequencing data and assessing whether any PRS associations were impacted

PRSs can use SNPs that do not achieve genome-wide significant p-value values in the original GWAS. This is of relevance as the GWAS p-value cutoff at which the PRS is most predictive is often unknown (21, 22). Following standard practice, we created PRSs using a predefined set of fixed GWAS p-value cutoffs (all SNPs with GWAS p-value < 1e-8, < 1e-7, < 1e-5, < 1e-3, < 1e-2 and < 1e-1), with each score distinguished by the number of SNPs used (**Fig. S3**). We then asked which of these PRSs were associated with occurrence with a phenotype of interest, accounting for multiple testing at a preselected false discovery rate of 10%.

**Rationale for Use of P+T versus Beta-Shrinkage**

We note that approaches that use beta shrinkage such as LDpred (23) have shown promise compared to pruning and thresholding (P+T). However, LDpred does not escape the issue of multiple testing due to the requirement that the fraction of causal markers be specified in advance of LDpred's beta shrinkage procedure. In the same way the p-value threshold impacts P+T, the fraction of causal markers parameter will impact the PRS produced by LDpred and relates to the genetic architecture of the trait. LDpred's fraction of causal variants parameter is less interpretable than P+T p-value threshold. LDpred also suffers from convergence issues due to the complexity of its fitting algorithm an issue observed by authors of a recent improvement on LDpred (see https://www.biorxiv.org/content/10.1101/375337v1). To our knowledge only one method GCTB (https://www.biorxiv.org/content/10.1101/522961v3) truly addresses the issue of multiple testing, but makes strong assumptions about the underlying LD data. We note that, because of its robustness, P+T is the benchmark to which methods are compared. We also point out that cross validation may be used instead of multiple testing correction to adjust the parameters of P+T, as well as LDpred, but requires a larger cohort than used in our study. Based on the considerations above, we opted for an older but more appropriate methodological approach, P+T, combined with FDR correction despite these stated shortcomings.

**PRS Associations with Skin irAE Occurrence**

We tested for associations between PRSs at a range of GWAS p-value cutoffs and occurrence of skin irAEs using logistic regression. False discovery rate (FDR) was estimated using the Benjamini-Hochberg (BH) procedure (24). Our analysis controlled for 5 genotype eigenvectors/PCs and gender. Logistic regression was performed using glm() in R (v3.5.0).

**PRS Associations with Survival and PRS by Trial Arm Interactions**

We tested for associations between PRSs at a range of GWAS p-value cutoffs and survival using a Cox proportional hazards model. p-values were computed using the Wald test on the coefficient associated with PRS. To identify trial arm by PRS interactions, p-values were computed using the Wald test on the coefficient associated with PRS by trial arm interaction term. The model for assessing trial arm interactions also contained the lower order terms for PRS and arm. False discovery rate (FDR) was estimated using the Benjamini-Hochberg (BH) procedure (24). Our analysis controlled for the following covariates, which were measured at baseline prior to treatment:

- Presence or absence of liver metastases.
- High or low CRP where (>10 mg/ml is high)
- Neutrophil to lymphocyte ratio normalized to the quantiles of the standard normal distribution.
- High or low alkaline phosphatase in serum at baseline (>147 IU/L was designated as high).
- High or low albumin in serum at baseline (<35g/L is low).
- High or low LDH in serum at baseline (>400 g/L is high).
- Gender
- Baseline ECOG status (0 or 1).
- 5 genotype PCs/eigenvectors

The same approach was used for tumor factors (TMB, T-effector signature, IC IHC, and TC IHC). For immune cell (IC) and tumor cell (TC) IHC of PD-L1 the integers 0,1,2,3 were used to capture the ordinal relationship between PD-L1 staining values. All survival analyses were conducted using the survival package in R (https://cran.r-project.org/web/packages/survival/index.html).

**Inference of HLA alleles from whole-genome sequencing data**

We used HLA*PRG:LA (retrieved March 8, 2017, git commit SHA-1 hash prefixed by 7b9ba45) to infer HLA alleles at G group resolution from whole-genome sequencing data, starting from BAM files generated as described above (25). HLA G groups are defined at 6-digit resolution, containing alleles with identical nucleotide sequence in the exons encoding the peptide binding domains. While it is possible that a G group contains a 6-digit allele with differences in its amino acid structure outside the binding domain, these alleles are very rare, and a G group is dominated by its common, eponymous allele. For example, out of 5 randomly selected non-C*06:02 members of the C:06:02:01 G group

(C*06:46N, C*06:55, C*06:73, C*06:236, C*06:262), only C*06:73 can be found at all across published data sets (www.allelefrequencies.net), with an allele frequency of 0.0001 in a Czech population of >5,000 individuals. Although HLA typing tools for WGS data have been consistently shown to be very accurate, more so than software inferring HLA alleles from SNP chip data, we performed in-house comparisons of HLA*PRG:LA, applied to our 30x WGS data, with Labcorp CLIA certified "gold standard" typing results in 56 individuals, achieving accuracies of 99.4% for class I, and 99.7% for class II genes.

**HLA associations with overall survival, and addition of HLA effects to disease-specific PRS**

We tested for associations between single HLA variants and overall survival (OS) using a Cox proportional hazards model, using the same set of covariates described above for PRS association testing. HLA associations are usually not considered in the selection of SNPs for polygenic risk scores, both due to their disproportionate contribution in terms of variance explained in many diseases, and the complexity of the MHC locus in terms of variability and linkage equilibrium. To assess whether inclusion of HLA variants can improve the PRS associations, we added the effect of HLA alleles previously reported to be associated with Psoriasis, Atopic Dermatitis, and Vitiligo to the respective PRS, making the following simplifying assumptions:

- Since HLA alleles were imputed at G group resolution, we assumed that all carriers of a given G group allele were carriers of the 4-digit allele of interest in the given G group.
- For reported associations of HLA amino acid residues, we identified carriers of those residues by the amino acid sequences of their inferred HLA alleles and categorized them accordingly (see Table S3). For reported associations on the level of HLA serotypes, we also categorized patients according to their carrier status for an allele belonging to the respective serotype (see Table S4).
- Given the strong associations of the HLA variants in the original studies (Table S3), we assumed they would reach the most stringent PRS p-value cutoff in the respective GWAS, and therefore added their effects to all PRS cutoffs.

HLA log odds ratios for risk were added to the PRS before quantile normalization, treating each allele as a further single variant adding to the polygenic model. Hence, we added the log odds ratio of the reported HLA allele associations times the number of copies of the allele to the PRS, followed by quantile normalization. We then performed association testing with overall survival and PRS+HLA by Trial Arm Interactions, using the same methodology as described above for the PRS analyses.

**Pre-Treatment Tumor RNA-seq**

The pathologic diagnosis of each case was confirmed by review of hematoxylin and eosin (H&E) stained slides and all samples that advanced to nucleic acid extraction contained a minimum of 20% tumor cells. H&E images were marked for macro-dissection by a pathologist. RNA (High Pure FFPET RNA Isolation Kit, Roche) was then extracted from the macro-dissected sections. Tumor RNA-seq data was generated using TruSeq RNA Access technology (Illumina). Reads were first aligned to ribosomal RNA sequences to remove ribosomal reads. The remaining reads were aligned to the human reference genome (NCBI Build 38) using GSNAP (26) version 2013-10-10, allowing maximum of two mismatches per 75 base sequence. (parameters: -M 2 -n 10 -B 2 -i 1 -N 1 -w 200000 -E 1 --pairmax-rna=200000 --clip-overlap).

To quantify gene expression levels, the number of reads mapped to the exons of each RefSeq gene was calculated using the functionality provided by the R package GenomicAlignments (Bioconductor) (27). RNA-seq library size L was estimated using the TMM method to estimate norm factors in edgeR (e.g. L = normFactor * total number of reads in sample). For each gene/sample count, we computed counts per million using a pseudo count (C + 0.5)/(L + 1) * 1e+06 following limma/voom where C was the read count of the gene in a given sample (28). The resulting counts per million values were scaled by gene length to obtain normalized RPKM values. We estimated 10 PEER factors directly on the normalized $\log_2$ counts per million values to account for batch effects in tumor RNA-seq (29).

**$CD8^+$ T-effector Signature Score Computation**

To construct the T-effector signature we used the following gene set of G=8 genes: CD8A, GZMA, GZMB, INFG, CXCL9, CXCL10, PRF1 and TBX21, as defined in (13). The signature was constructed by first z transforming the $\log_2$ counts per million values for each gene in the gene set across the N tumor RNA seq samples in a given data set and then performing a singular value decomposition (principal component analysis) of the resulting G x N data matrix. The first principal component (PC1) is by definition the T-effector signature for the given data set. It is by construction a weighted sum of the genes in the gene set, focusing the score on the largest block of well-correlated genes in the set, while down-weighing contributions from genes that do not track with other members of the gene set.

**Additional Tumor Factors**

Data on immune cell (IC) and tumor cell (TC) staining of PD-L1 by IHC, as well as tumor mutation burden, was obtained using methods described in the previous publication and protocols for IMvigor211 (14). xCell cell type enrichment scores were obtained by entering the normalized RPKM values at the xCell website http://xcell.ucsf.edu (30).

**Assessing Relevance of PRS in Differing Tumor Immune Contexts**

We statistically assessed whether combining a PRS and a tumor factor was more informative of treatment effect on overall survival than a PRS alone. We focused on tumor gene expression of T-cell differentiation, recruitment and response cytokines and chemokines, filtering for those genes with median RPKM >0.1 and median read count of >10 across individuals (**Table S5**). We limited our analysis the most strongly predictive PRSs of overall survival as reflected by the p-value of the PRS by trial arm interaction and identified by their GWAS p-value cutoff (**Fig. 3a**). PRSs and tumor gene expression values were transformed into categorical variables (high and low) on the basis of median splits across the entire IMvigor211 population. This approach made minimal assumptions about the relative scale of PRSs and tumor gene expression values. We tested for a non-zero 3-way interaction term between PRS (high/low) by trial arm (atezolizumab/chemotherapy) by tumor factor/RNA-seq (high/low) in a Cox proportional hazards model for overall survival. The Cox proportional hazards model also contained all lower order interaction terms. p-values were computed using the Wald test on the coefficient of this 3-way interaction term using the survival package in R. False discovery rate (FDR) was estimated using the Benjamini-Hochberg (BH) procedure (24). We controlled for the same baseline factors above in our tests for association between PRSs and survival. We additionally included 10 PEER factors to account for batch effects in tumor RNA-seq data. Formally, the underlying statistical test rejected the null hypothesis that the difference between the hazard ratios at high or low PRS was the same in individuals with high or low tumor factor value.

**Table S1.** GWAS abbreviations, citations, and summary statistic URLs

| Abbreviation | GWAS | Cases/Controls | Citation | URLs |
|---|---|---|---|---|
| AD | Atopic dermatitis | 10788/37217 | (31) | https://data.bris.ac.uk/data/dataset/28uchsdpmub118uex26ylacqm |
| PSO/IC | Psoriasis Immunochip | 2997/9183 | (32) | https://www.immunobase.org/downloads/protected_data/iChip_Data ftp://ftp.ebi.ac.uk/pub/databases/gwas/summary_statistics/TsoiLC_23143594_GCST005527 |
| PSO/UKBB | Self-Reported Psoriasis UK Biobank | 3871/333288 | (33) | http://www.nealelab.is/uk-biobank (round 1 results) |
| VIT | Vitiligo | 4680/39586 | (34) | ftp://ftp.ebi.ac.uk/pub/databases/gwas/summary_statistics/JinY_27723757_GCST004785 |
| ALZ | Alzheimer's disease | 17008/37154 | (35) | http://web.pasteur-lille.fr/en/recherche/u744/igap/igap_download.php |

| | | | Skin irAE | | | OS atezo | | | OS chemo | | | OS interaction | | |
|---|---|---|---|---|---|---|---|---|---|---|---|---|---|---|
| gwas.cur | pcutoff | N.snps | beta (95% CI) | *p.val* | *p.adj* | beta (95% CI) | *p.val* | *p.adj* | beta (95% CI) | *p.val* | *p.adj* | beta (95% CI) | *p.val* | *p.adj* |
| **PSO/IC** | 1.00E-08 | 61 | **0.58 (0.22 — 0.94)** | **0.0016** | **0.04** | -0.11 (-0.29 — 0.06) | 0.21 | 0.31 | 0.16 (-0.01 — 0.32) | 0.07 | 0.42 | **0.23 (0.00 — 0.47)** | **0.052** | **0.10** |
| | 1.00E-07 | 76 | **0.52 (0.17 — 0.87)** | **0.0038** | **0.05** | -0.13 (-0.31 — 0.04) | 0.14 | 0.22 | 0.14 (-0.02 — 0.31) | 0.10 | 0.42 | **0.23 (0.00 — 0.47)** | **0.050** | **0.10** |
| | 1.00E-05 | 143 | **0.41 (0.05 — 0.77)** | **0.025** | **0.10** | -0.15 (-0.33 — 0.03) | 0.10 | 0.18 | 0.11 (-0.06 — 0.27) | 0.21 | 0.43 | **0.24 (0.01 — 0.47)** | **0.045** | **0.10** |
| | 0.001 | 493 | 0.27 (-0.07 — 0.60) | 0.12 | 0.35 | **-0.21 (-0.38 — -0.03)** | **0.019** | **0.07** | 0.11 (-0.06 — 0.29) | 0.19 | 0.43 | **0.29 (0.05 — 0.52)** | **0.016** | **0.06** |
| | 0.01 | 1396 | 0.24 (-0.10 — 0.58) | 0.17 | 0.40 | **-0.25 (-0.43 — -0.07)** | **0.006** | **0.07** | 0.12 (-0.05 — 0.30) | 0.15 | 0.43 | **0.33 (0.09 — 0.56)** | **0.006** | **0.03** |
| | 0.1 | 6033 | 0.08 (-0.27 — 0.42) | 0.67 | 0.76 | **-0.24 (-0.41 — -0.06)** | **0.008** | **0.07** | 0.23 (0.04 — 0.42) | 0.02 | 0.42 | **0.39 (0.15 — 0.62)** | **0.0011** | **0.02** |
| **PSO/UKBB** | 1.00E-08 | 45 | 0.35 (-0.01 — 0.71) | 0.06 | 0.20 | **-0.19 (-0.36 — -0.01)** | **0.038** | **0.10** | 0.10 (-0.06 — 0.27) | 0.23 | 0.43 | **0.29 (0.06 — 0.53)** | **0.016** | **0.06** |
| | 1.00E-07 | 69 | **0.44 (0.07 — 0.81)** | **0.02** | **0.10** | **-0.24 (-0.42 — -0.06)** | **0.010** | **0.07** | 0.14 (-0.03 — 0.30) | 0.10 | 0.42 | **0.38 (0.14 — 0.61)** | **0.0021** | **0.02** |
| | 1.00E-05 | 217 | **0.40 (0.05 — 0.75)** | **0.02** | **0.10** | **-0.19 (-0.36 — -0.03)** | **0.021** | **0.07** | 0.05 (-0.12 — 0.21) | 0.58 | 0.73 | **0.24 (0.01 — 0.47)** | **0.045** | **0.10** |
| | 0.001 | 3927 | **0.45 (0.10 — 0.81)** | **0.01** | **0.09** | -0.09 (-0.25 — 0.08) | 0.31 | 0.44 | 0.08 (-0.09 — 0.25) | 0.34 | 0.53 | 0.14 (-0.09 — 0.37) | 0.24 | 0.35 |
| | 0.01 | 26981 | 0.21 (-0.11 — 0.53) | 0.20 | 0.40 | -0.03 (-0.19 — 0.12) | 0.68 | 0.73 | -0.03 (-0.19 — 0.14) | 0.74 | 0.83 | 0.00 (-0.22 — 0.22) | 0.99 | 0.99 |
| | 0.1 | 215596 | 0.12 (-0.18 — 0.41) | 0.44 | 0.62 | -0.06 (-0.20 — 0.08) | 0.37 | 0.49 | -0.17 (-0.37 — 0.03) | 0.09 | 0.42 | -0.11 (-0.35 — 0.13) | 0.37 | 0.44 |
| **VIT** | 1.00E-08 | 79 | -0.10 (-0.44 — 0.24) | 0.56 | 0.68 | **-0.19 (-0.37 — -0.01)** | **0.035** | **0.10** | 0.14 (-0.04 — 0.31) | 0.13 | 0.43 | **0.35 (0.11 — 0.58)** | **0.0037** | **0.02** |
| | 1.00E-07 | 100 | -0.04 (-0.39 — 0.31) | 0.81 | 0.81 | **-0.21 (-0.39 — -0.03)** | **0.023** | **0.07** | 0.12 (-0.05 — 0.30) | 0.17 | 0.43 | **0.35 (0.11 — 0.58)** | **0.0036** | **0.02** |
| | 1.00E-05 | 282 | -0.15 (-0.51 — 0.22) | 0.44 | 0.62 | **-0.22 (-0.41 — -0.03)** | **0.023** | **0.07** | 0.06 (-0.11 — 0.23) | 0.47 | 0.63 | **0.28 (0.04 — 0.52)** | **0.021** | **0.06** |
| | 0.001 | 3008 | -0.19 (-0.68 — 0.29) | 0.44 | 0.62 | -0.21 (-0.46 — 0.05) | 0.11 | 0.19 | 0.11 (-0.09 — 0.31) | 0.30 | 0.52 | 0.21 (-0.02 — 0.44) | 0.079 | 0.13 |
| | 0.01 | 16974 | 0.20 (-0.41 — 0.81) | 0.52 | 0.68 | -0.31 (-0.63 — 0.01) | 0.056 | 0.13 | 0.29 (0.01 — 0.56) | 0.04 | 0.42 | 0.22 (-0.01 — 0.45) | 0.059 | 0.10 |
| | 0.1 | 103016 | -0.19 (-0.84 — 0.46) | 0.57 | 0.68 | -0.32 (-0.67 — 0.03) | 0.073 | 0.15 | 0.19 (-0.10 — 0.49) | 0.20 | 0.43 | 0.17 (-0.05 — 0.39) | 0.12 | 0.19 |
| **AD** | 1.00E-08 | 20 | -0.23 (-0.57 — 0.11) | 0.19 | 0.40 | **0.21 (0.04 — 0.38)** | **0.018** | **0.07** | -0.14 (-0.30 — 0.01) | 0.07 | 0.42 | **-0.35 (-0.58 — -0.13)** | **0.0020** | **0.02** |
| | 1.00E-07 | 33 | -0.18 (-0.50 — 0.15) | 0.29 | 0.50 | 0.14 (-0.03 — 0.31) | 0.11 | 0.19 | -0.10 (-0.26 — 0.06) | 0.22 | 0.43 | **-0.25 (-0.48 — -0.02)** | **0.032** | **0.08** |
| | 1.00E-05 | 101 | -0.22 (-0.56 — 0.12) | 0.20 | 0.40 | 0.16 (-0.01 — 0.33) | 0.066 | 0.14 | -0.07 (-0.24 — 0.10) | 0.40 | 0.58 | **-0.26 (-0.49 — -0.03)** | **0.030** | **0.08** |
| | 0.001 | 2796 | -0.07 (-0.45 — 0.31) | 0.72 | 0.77 | 0.06 (-0.13 — 0.24) | 0.56 | 0.71 | -0.09 (-0.28 — 0.09) | 0.31 | 0.52 | -0.11 (-0.34 — 0.11) | 0.33 | 0.42 |
| | 0.01 | 21027 | -0.06 (-0.44 — 0.31) | 0.74 | 0.77 | -0.05 (-0.23 — 0.14) | 0.62 | 0.73 | -0.01 (-0.21 — 0.19) | 0.93 | 0.96 | 0.05 (-0.18 — 0.29) | 0.65 | 0.70 |
| | 0.1 | 159100 | 0.27 (-0.16 — 0.70) | 0.22 | 0.41 | -0.06 (-0.27 — 0.16) | 0.62 | 0.73 | 0.04 (-0.16 — 0.24) | 0.71 | 0.83 | 0.06 (-0.17 — 0.30) | 0.60 | 0.67 |
| **ALZ** | 1.00E-08 | 66 | -0.13 (-0.46 — 0.19) | 0.41 | NA | -0.03 (-0.21 — 0.14) | 0.70 | NA | 0.00 (-0.18 — 0.18) | 0.97 | NA | 0.03 (-0.21 — 0.27) | 0.83 | NA |
| | 1.00E-07 | 76 | -0.17 (-0.49 — 0.15) | 0.30 | NA | -0.01 (-0.18 — 0.17) | 0.95 | NA | 0.00 (-0.18 — 0.17) | 0.96 | NA | 0.00 (-0.23 — 0.24) | 0.99 | NA |
| | 1.00E-05 | 146 | -0.12 (-0.44 — 0.20) | 0.46 | NA | -0.05 (-0.22 — 0.12) | 0.58 | NA | 0.01 (-0.17 — 0.19) | 0.91 | NA | 0.04 (-0.20 — 0.28) | 0.73 | NA |
| | 0.001 | 1931 | -0.03 (-0.35 — 0.28) | 0.83 | NA | -0.02 (-0.18 — 0.14) | 0.78 | NA | -0.02 (-0.19 — 0.14) | 0.81 | NA | 0.00 (-0.23 — 0.22) | 0.98 | NA |
| | 0.01 | 12488 | -0.07 (-0.42 — 0.28) | 0.69 | NA | 0.02 (-0.15 — 0.19) | 0.79 | NA | -0.14 (-0.31 — 0.04) | 0.13 | NA | -0.16 (-0.38 — 0.07) | 0.18 | NA |
| | 0.1 | 85131 | 0.01 (-0.35 — 0.36) | 0.97 | NA | 0.06 (-0.11 — 0.24) | 0.48 | NA | -0.12 (-0.31 — 0.07) | 0.22 | NA | -0.18 (-0.40 — 0.05) | 0.13 | NA |
| **TUMOR** | T-eff | NA | - | - | - | **-0.22 (-0.39 — -0.04)** | **0.014** | **0.07** | -0.07 (-0.25 — 0.10) | 0.41 | 0.58 | 0.05 (-0.04 — 0.15) | 0.27 | 0.37 |
| | TMB | NA | - | - | - | -0.03 (-0.28 — 0.21) | 0.78 | 0.81 | -0.02 (-0.25 — 0.20) | 0.84 | 0.90 | 0.02 (-0.02 — 0.06) | 0.31 | 0.42 |
| | IC IHC | NA | - | - | - | -0.04 (-0.46 — 0.37) | 0.85 | 0.85 | -0.01 (-0.43 — 0.41) | 0.96 | 0.96 | 0.02 (-0.56 — 0.59) | 0.96 | 0.99 |
| | TC IHC | NA | - | - | - | -0.11 (-0.58 — 0.36) | 0.65 | 0.73 | 0.10 (-0.37 — 0.58) | 0.67 | 0.82 | 0.29 (-0.35 — 0.94) | 0.38 | 0.44 |

**Table S2.** Provided as an Excel .XLSX file. Tabs correspond to phenotypes associated with the PRSs. Columns are defined as follows.

| **Column** | **Description** |
|---|---|
| gwas.cur | GWAS from which PRS was derived |
| p.cutoff | GWAS p-value cutoff used to construct PRS |
| N.snps | number of variants in the PRS |
| beta | coefficient associated with PRS term or interaction beta for PRS by arm interaction test |
| low.95, high.95 | 95% confidence interval around PRS/interaction coefficient |
| p.val | p-value of statistical test of association or interaction |
| p.adj | adjusted p-value by the Benjamini Hochberg procedure |

**Table S3.** Selection of HLA alleles for statistical testing

| Allele, Amino Acid residue or Serotype | Disease | P | OR | 4-digit alleles covered by AA position or serotype |
|---|---|---|---|---|
| HLA-C*06:02 | Psoriasis | 2.1E-201 | 3.26 | |
| HLA-C*12:03 | Psoriasis | 6.5E-12 | 1.38 | |
| HLA-B p.67 Cys | Psoriasis | 6.0E-35 | 1.56 | B*14:02,B*15:10,B*15:18,B*27:02, B*27:05,B*39:01 |
| HLA-B p.67 Met | Psoriasis | 2.6E-13 | 1.44 | B*57:01,B*57:03,B*58:02 |
| HLA-B p.9 Asp | Psoriasis | 1.6E-09 | 1.33 | B*08:01 |
| HLA-A p.95 Val | Psoriasis | 4.7E-28 | 1.31 | A*02:01,A*02:06,A*69:01 |
| HLA-DRB1*07:01 | Atopic Dermatitis | 1.4E-07 | 0.65 | |
| B*44:02 | Atopic Dermatitis | 9.6E-05 | 1.39 | |
| HLA-A02 | Vitiligo | <0.0001 | 1.52 | A*02:01,A*02:02,A*02:03, A*02:05,A*02:06,A*02:07,A*02:11 |
| HLA-A33 | Vitiligo | <0.0001 | 2.23 | A*33:01, A*33:03 |

HLA alleles, amino acid residues, and serotypes were selected based on published associations with psoriasis (36), atopic dermatitis (37), and vitiligo (38).

**Table S4.** HLA associations with OS in IMvigor211 atezolizumab and chemotherapy arms

| Allele, Amino Acid residue or Serotype | Atezolizumab Arm | | | Chemotherapy Arm | | |
|---|---|---|---|---|---|---|
| | HR | Std Error | P | HR | Std Error | P |
| HLA-B*44:02:01G | 0.98 | 0.28 | 0.93 | 1.08 | 0.26 | 0.76 |
| HLA-C*06:02:01G | 0.77 | 0.25 | 0.30 | 0.78 | 0.25 | 0.32 |
| HLA-C*12:03 | 0.86 | 0.26 | 0.57 | 1.04 | 0.25 | 0.89 |
| HLA-DRB1*07:01:01G | 0.98 | 0.19 | 0.90 | 0.86 | 0.20 | 0.43 |
| HLA-B p.67 Cys | 0.78 | 0.25 | 0.31 | 1.13 | 0.23 | 0.60 |
| HLA-B p.67 Met | 1.09 | 0.35 | 0.80 | 1.05 | 0.38 | 0.90 |
| HLA-B p.9 Asp | 2.20 | 0.43 | 0.07 | 0.87 | 0.36 | 0.69 |
| HLA-A p.95 Val | 1.11 | 0.17 | 0.54 | 1.15 | 0.17 | 0.42 |
| HLA-A02 | 1.14 | 0.17 | 0.45 | 1.17 | 0.17 | 0.35 |
| HLA-A33 | 0.77 | 0.47 | 0.58 | 1.49 | 0.50 | 0.42 |

Selected HLA alleles, amino acid residues, and serotypes were tested for association with overall survival in IMvigor211 atezolizumab and chemotherapy arms. Hazard ratio, HR < 1, indicates that carriers of the HLA allele have better overall survival.

**Table S5.** Genes Combined with PRSs to Define Patient Subgroups

| Gene Set Description | Genes |
|---|---|
| Treg Differentiation | IL2, TGFB1 |
| Treg Recruitment | CCL20 |
| Treg Response | TGFB1, IL10, *IL35* (EBI3, IL12A) |
| Th1 Differentiation | *IL12* (IL12A, IL12B) |
| Th1 Recruitment | CXCL9, CXCL10, CXCL11 |
| Th1 Response | IFNG, TNF |
| Th2 Differentiation | IL4 |
| Th2 Response | IL4, IL13 |
| Th9 Differentiation | IL4, TGFB1 |
| Th9 Response | IL9, IL10 |
| Th17 Differentiation | IL6, *IL23* (IL12A, IL23A), IL1B, TGFB1 |
| Th17 Recruitment | CCL20 |
| Th17 Response | IL17A, IL17F, CXCL1, CXCL2, CXCL5 |
| Th22 Differentiation | IL6, TNF |
| Th22 Response | IL22 |
| Tfh Differentiation | IL6, IL21 |
| Tfh Response | IL6, IL21 |
| $CD8^+$ T-effector | CD8A, GZMA, GZMB, IFNG, CXCL9, CXCL10, PRF1, TBX21 |

Cytokines that are heterodimers are italicized with genes in parentheses. Genes with median RPKM >0.1 and median count >10 across individuals in bulk tumor RNA-seq are highlighted in red. Gene sets were adapted from (3, 13, 39).

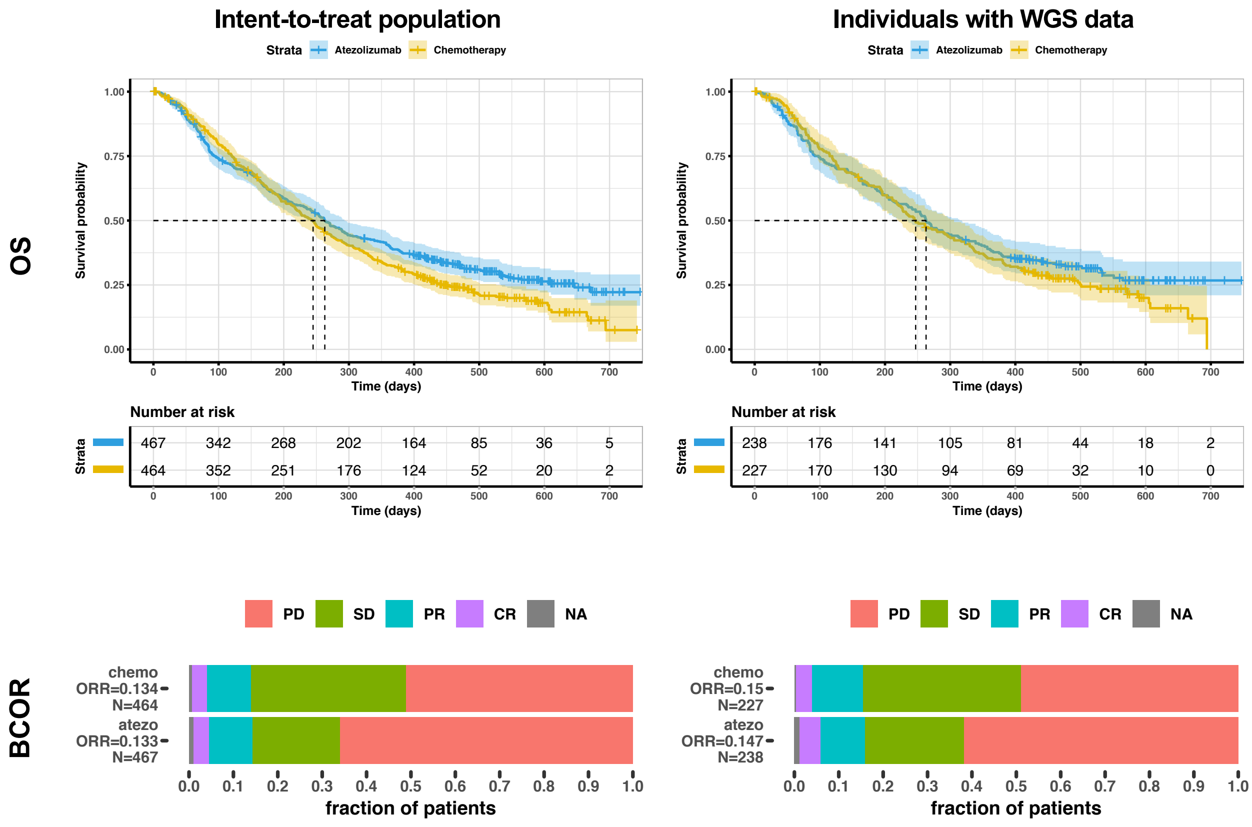


**Figure S1.** Comparison of outcomes in the intent-to-treat population (ITT) and the subpopulation with whole genome germline sequencing (WGS) data. The left column shows KM plots for overall survival (OS) and the distribution of Best Confirmed Objective Response (BCOR) in the intent-treat-population (ITT) of N=931 individuals. CR, PR, SD, and PD designate complete response, partial response, stable disease, and progressive disease respectively. Corresponding arm-specific objective response rates (ORRs) are shown to the left of the stacked proportional bar plots. The right column shows KM plots for OS and BCOR rates for N=465 with WGS data meeting strict criteria for population and genotype data quality control. By testing for an arm by population interaction, we confirmed that the hazard ratio between arms for OS (p=0.70) and the odds ratio between arms associated with response (p=0.98) were not statistically different.

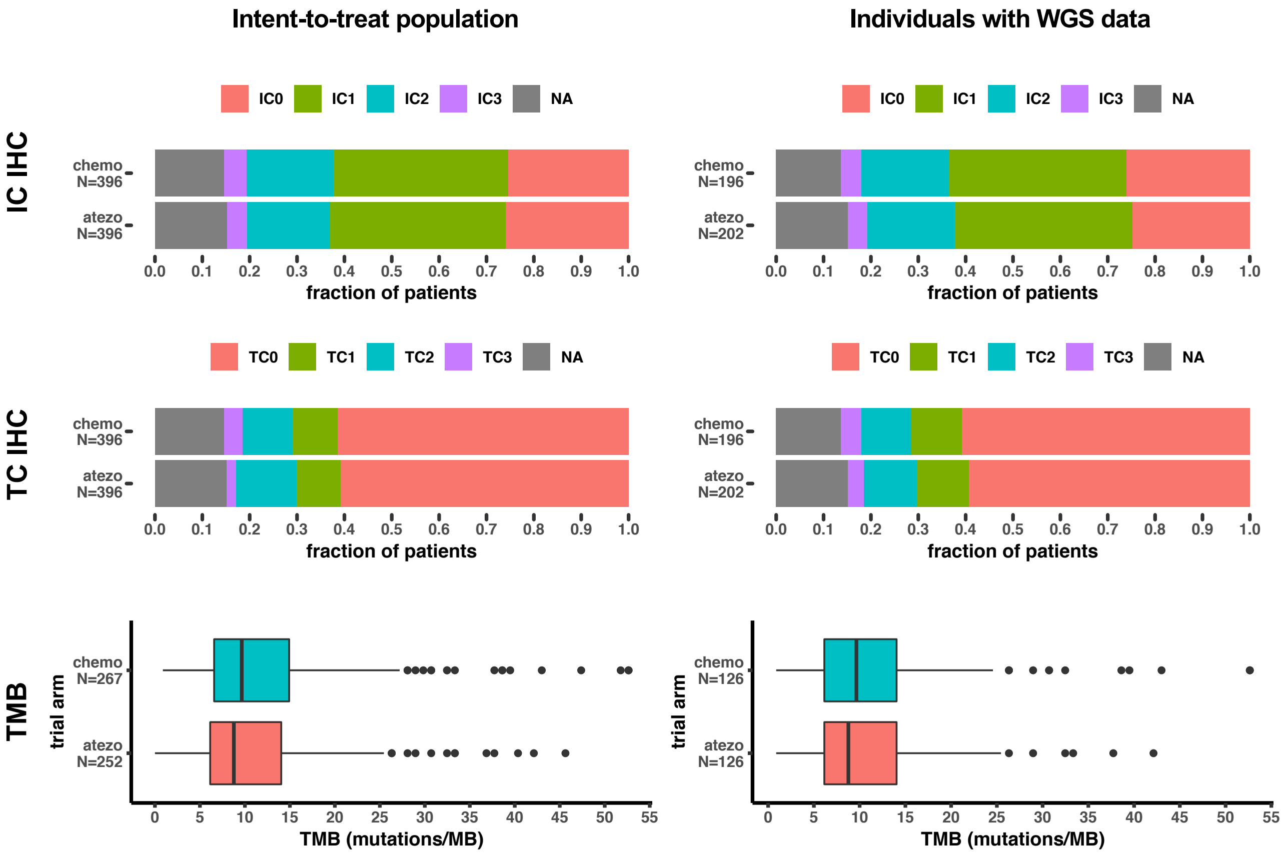


**Figure S2.** Comparison of tumor factors in the intent-to-treat population (ITT) and whole genome germline sequencing (WGS) available population. Here, N designates the number individuals for which the tumor factor was measured in the ITT and the subpopulation. NA designates the proportion of individuals for which the tumor factor was not measured. TC0 and IC0 designate tumor samples with no evidence of immune and tumor cell staining of PD-L1 by IHC respectively. TC1-TC3 and IC1-IC3 designate increasing levels of tumor and immune cell PD-L1 staining as previously defined (14). We confirmed that proportion of TC0/TC1 vs. TC2/3 individuals across arms (p=0.79) and IC0/IC1 vs. IC2/IC3 individuals across arms did not differ across the ITT or WGS available population. We additionally confirmed that the difference in the mean tumor mutation burden (TMB) across arms did not change significantly (p=0.80).

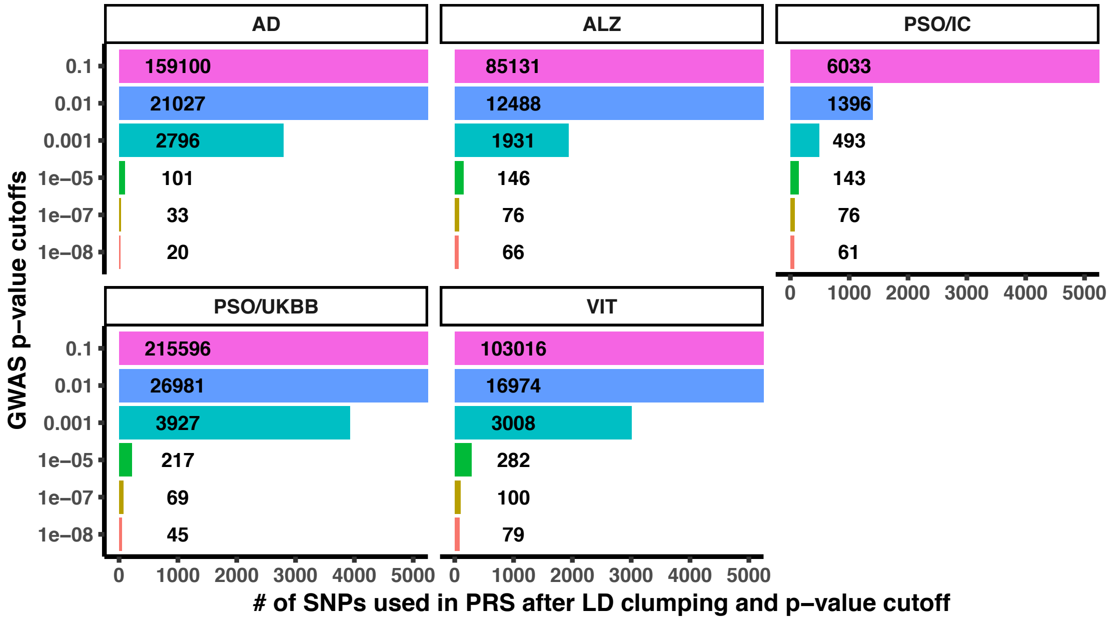


**Figure S3.** Number of SNPs at a given GWAS p-value cutoff. Bar plots were clipped at 5000 SNPs. GWAS p-value cutoff was applied after filtering and LD clumping. The total number of SNPs used per PRS and GWAS p-value cutoff is overlaid on the bars. See **Table 1** for details on the original GWAS studies used to construct PRSs and Methods for selecting these SNPs.

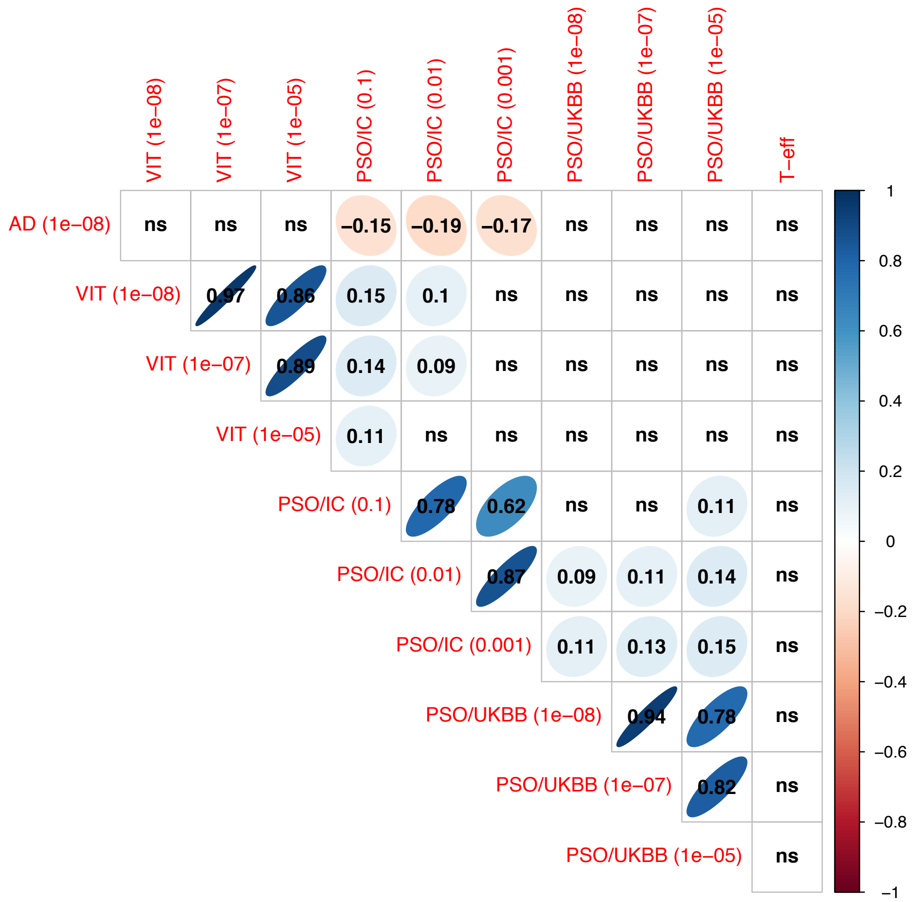


**Figure S4.** Spearman's rank correlation between PRSs and the T-effector signature across individuals in IMvigor211. Only PRSs significantly associated with OS in the atezolizumab arm at an FDR of 10% are shown. PRS GWAS p-value cutoffs are provided in parentheses. Rank correlations with $p \geq 0.05$ are labeled ns. As expected PRSs at differing GWAS p-value cutoffs were highly correlated. Notably the atopic dermatitis (AD) PRS was negatively correlated with psoriasis PRSs where cases were determined clinically (PSO/IC). A self-reported psoriasis GWAS PRSs (PSO/UKBB) only weakly correlated with PRSs derived for clinically determined psoriasis cases.

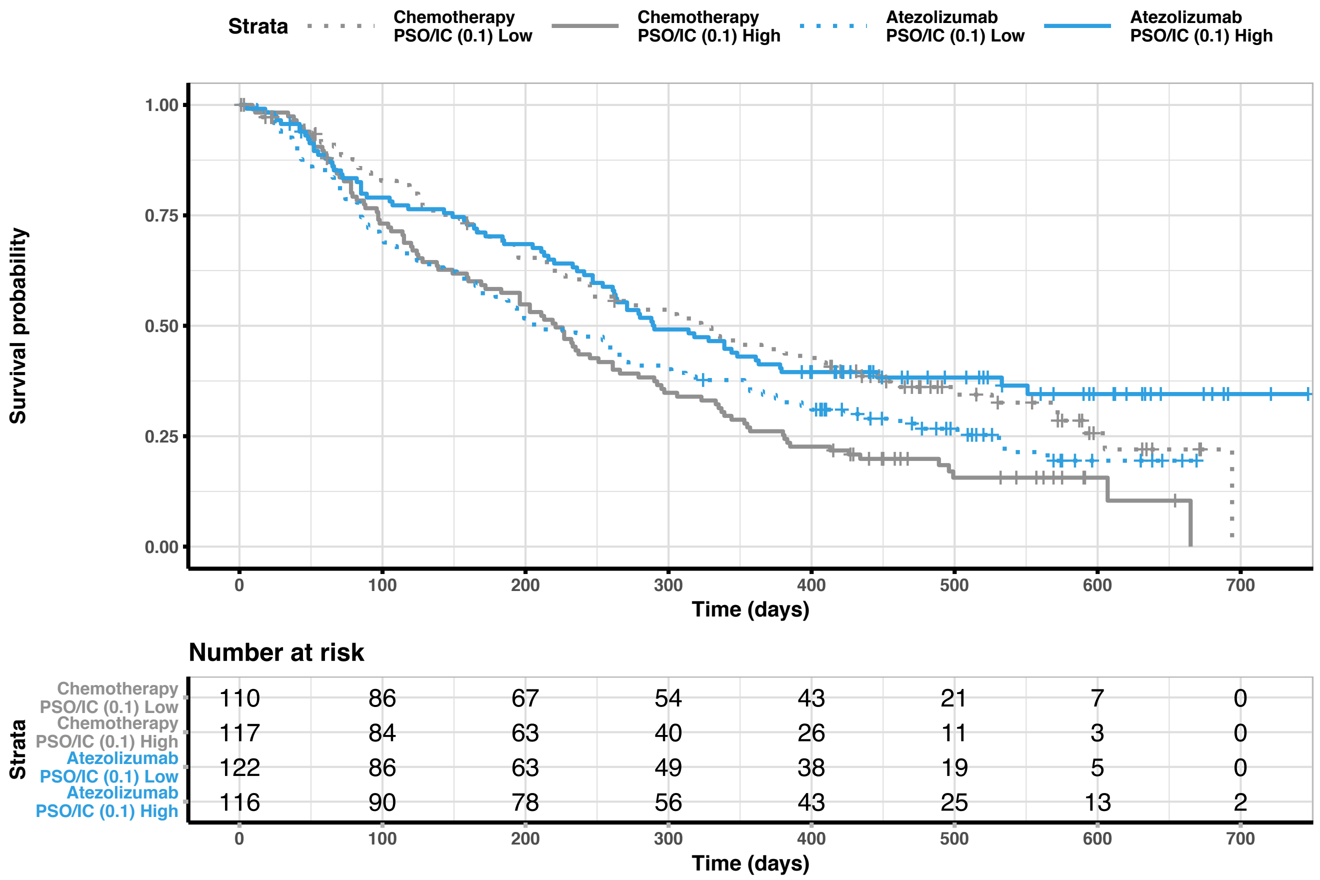


**Figure S5.** Kaplan-Meir plot of overall survival (OS) comparing atezolizumab to chemotherapy for individuals that had high or low genetic risk for psoriasis. Plot used the PRS derived from the PSO/IC psoriasis study at a GWAS p-value cutoff of 0.1. Atezolizumab OS data is plotted in blue. Dotted lines show the low risk group. Tick marks designate censoring.

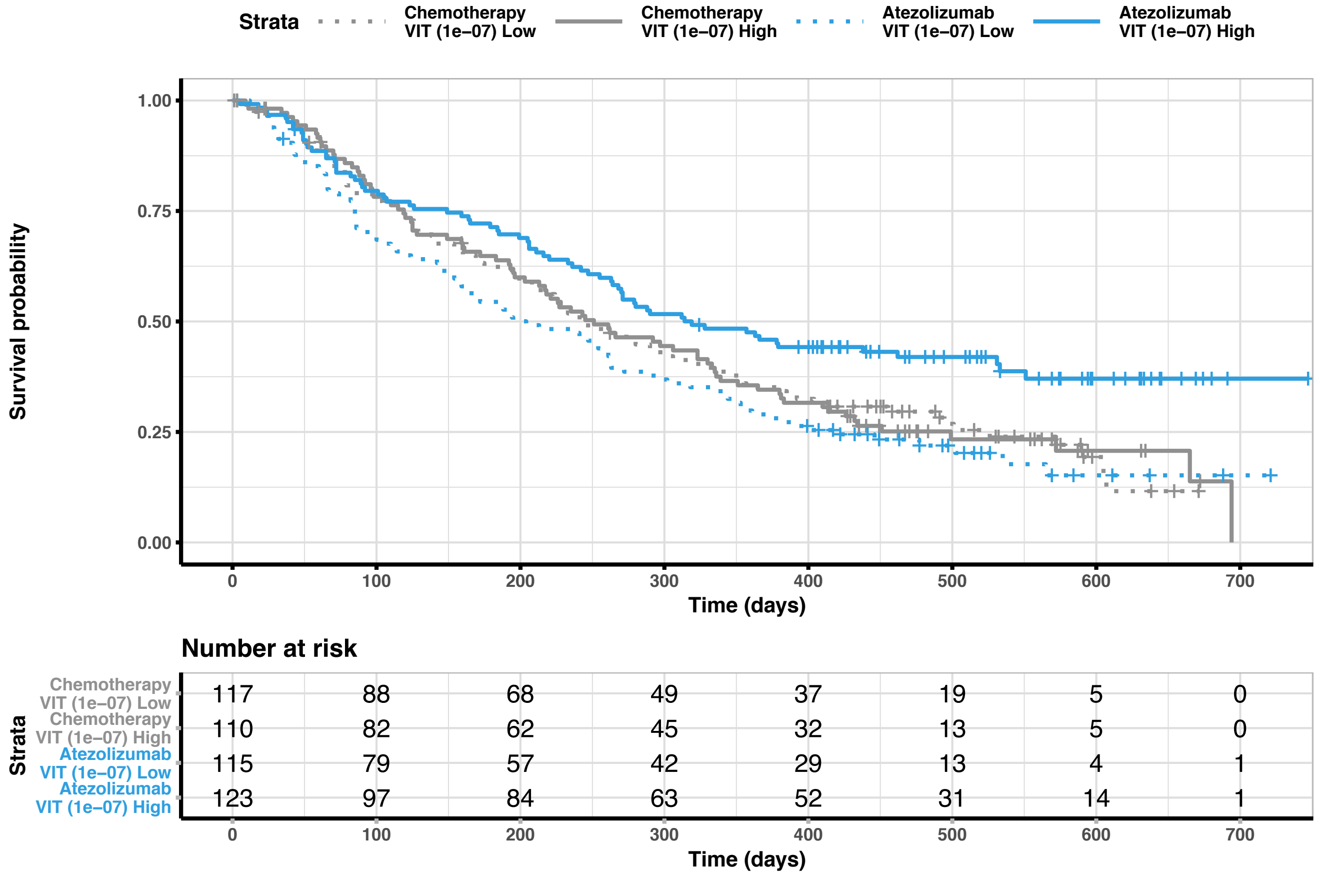


**Figure S6.** Kaplan-Meir plot of overall survival (OS) comparing atezolizumab to chemotherapy for individuals that had high or low genetic risk for vitiligo. Plot used the PRS derived from SNPs that met a GWAS p-value cutoff of 1e-07. Atezolizumab OS data is plotted in blue. Dotted lines show the low risk group. Tick marks designate censoring.

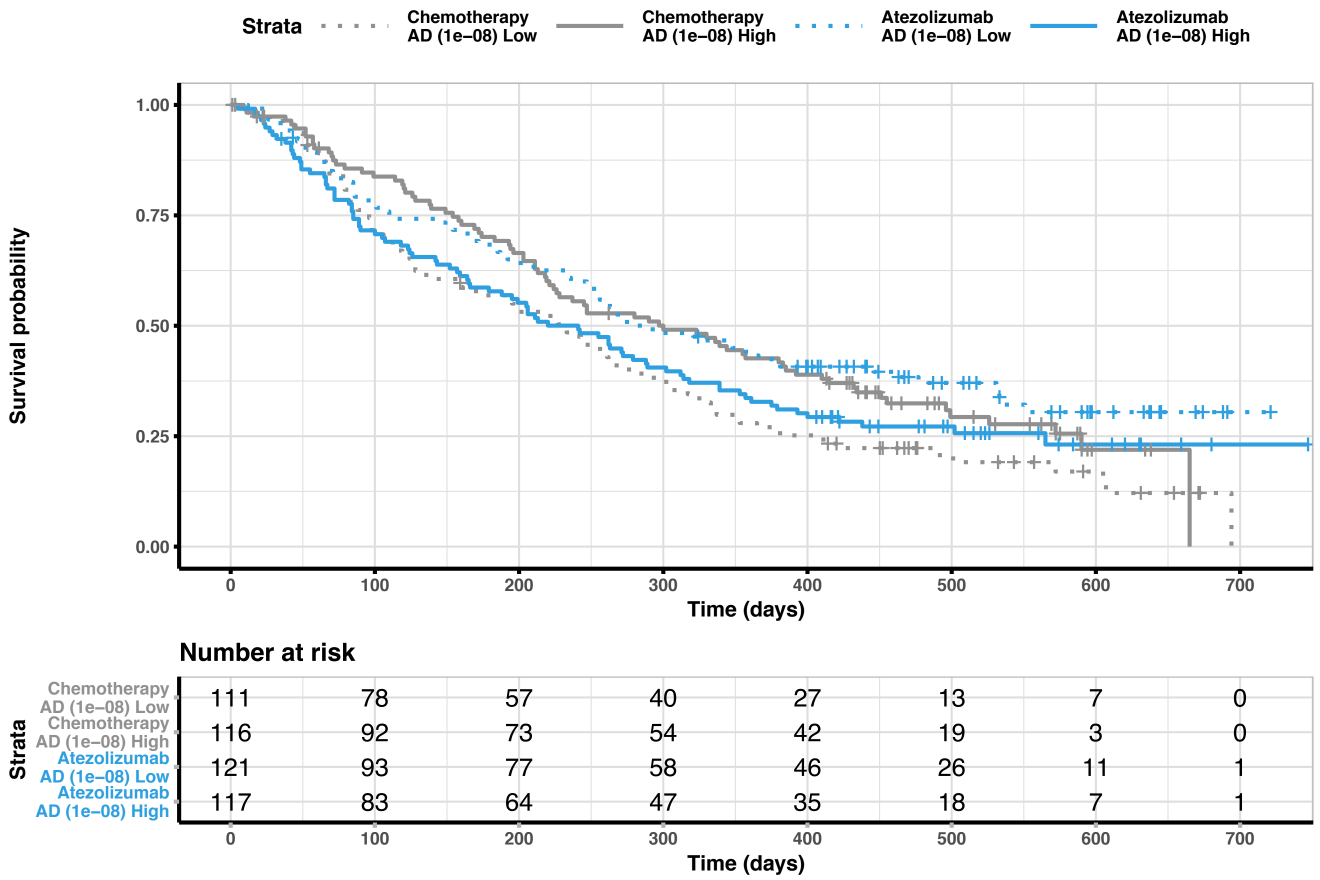


**Figure S7.** Kaplan-Meir plot of overall survival (OS) comparing atezolizumab to chemotherapy for individuals that had high or low genetic risk for atopic dermatitis (AD). Plot used the PRS derived from SNPs that met a GWAS p-value cutoff of 1e-08. Atezolizumab OS data is plotted in blue. Dotted lines show the low risk group. Tick marks designate censoring.

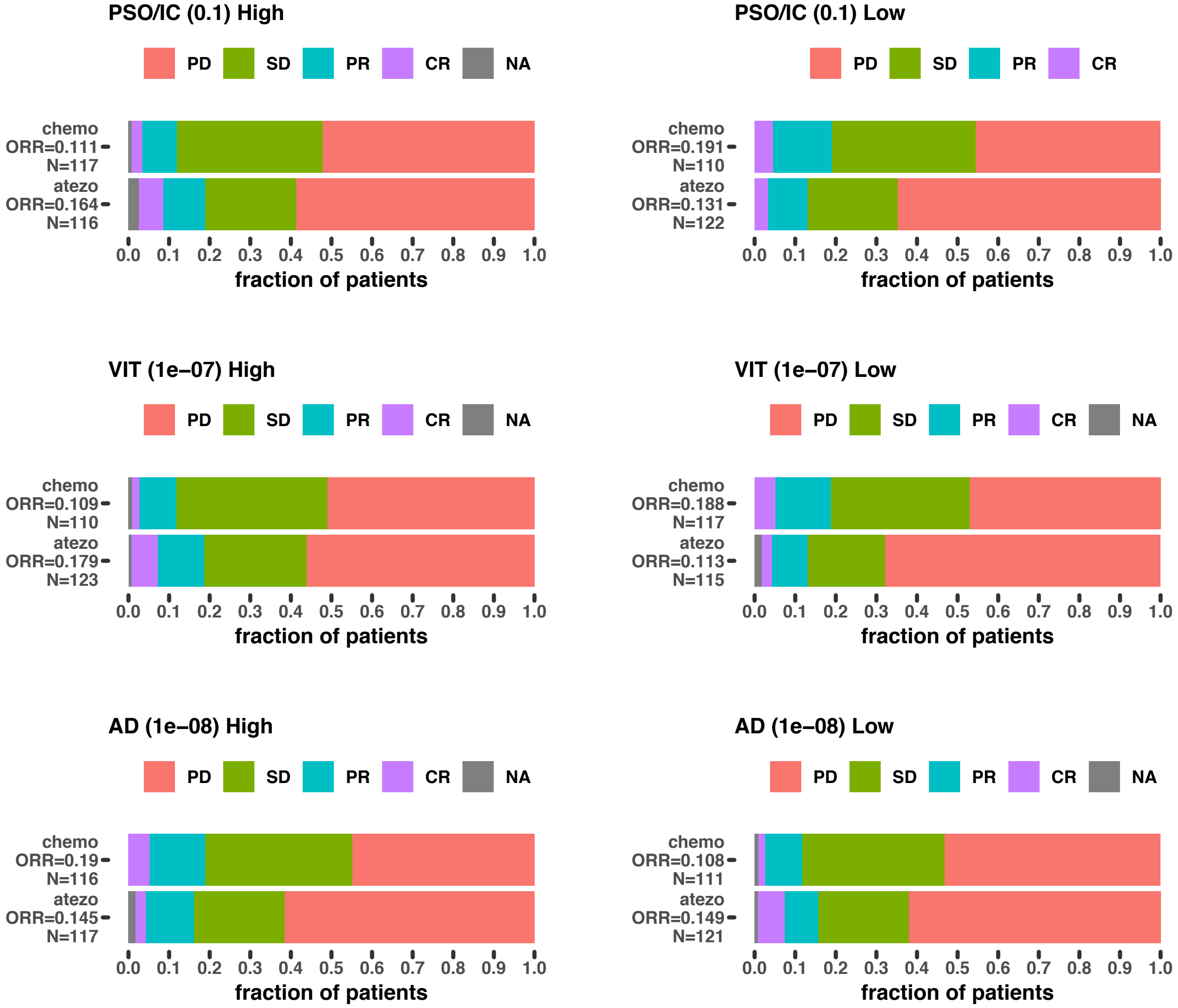


**Figure S8.** Objective response rates (ORRs) reflect patterns in OS KM plots. High and low genetic risk groups were defined using PRSs that were the most strongly predictive as measured by a PRS by trial arm interaction. The corresponding GWAS p-value cutoff used for the PRS is shown in parentheses. PD, SD, PR, and CR indicate progressive disease, stable disease, partial response, and complete response respectively. NA designates individuals where best confirmed object response (BCOR) data was unavailable.

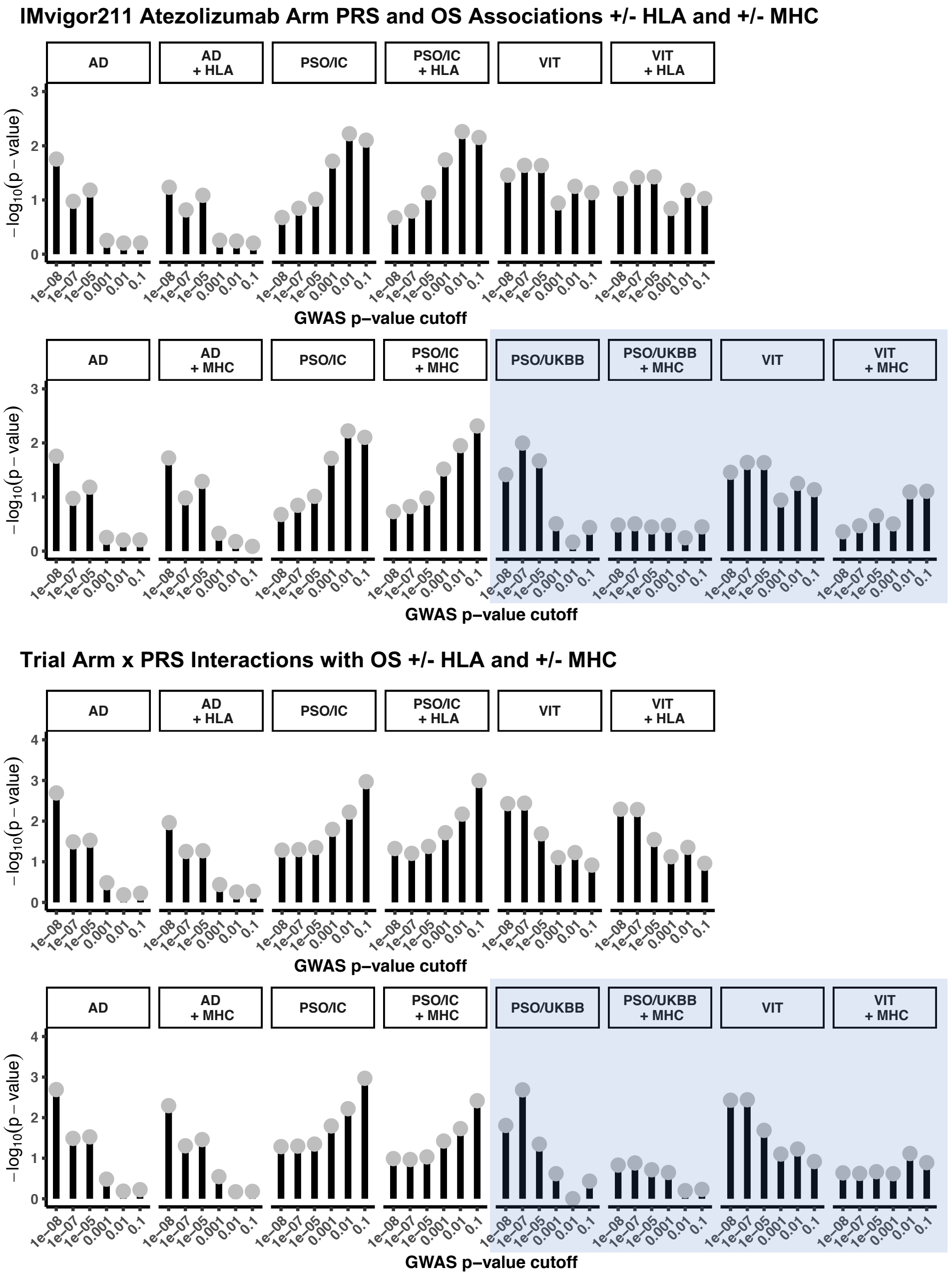


**Figure S9.** Negative $\log_{10}$ p-values for a given GWAS and p-value cutoff PRS with and without contribution of the MHC region or relevant HLA alleles, testing for association with overall survival in the atezolizumab arm (upper panel, related to Fig. 2d), and for a statistically significant trial arm by PRS interaction (lower panel, related to Fig. 3a), using a Cox proportional hazards model for overall survival (OS), controlling for 5 genotype principal components and several baseline clinical factors (see Methods). Light blue regions show PRSs where the association was attenuated by inclusion of variants from MHC region, likely due to a poor approximation of linkage disequilibrium. HLA allele effects could not be considered for the PSO/UKBB study, as the study used a linear mixed model with a response variable set to 1 for cases and -1 for controls to identify associations in a cohort with orders of magnitude more controls than cases (33). The coefficients provided in the summary statistics for the PSO/UKBB study were not directly interpretable as log odds ratios.

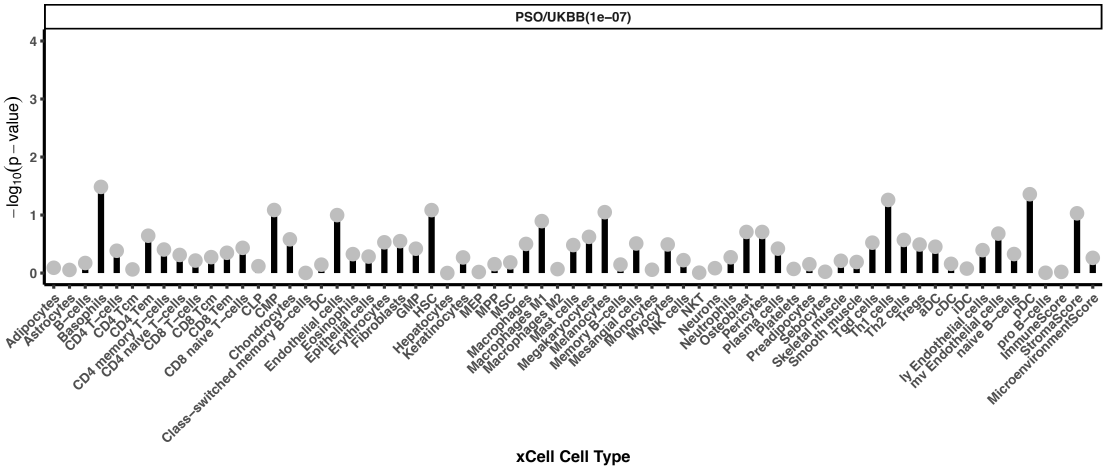


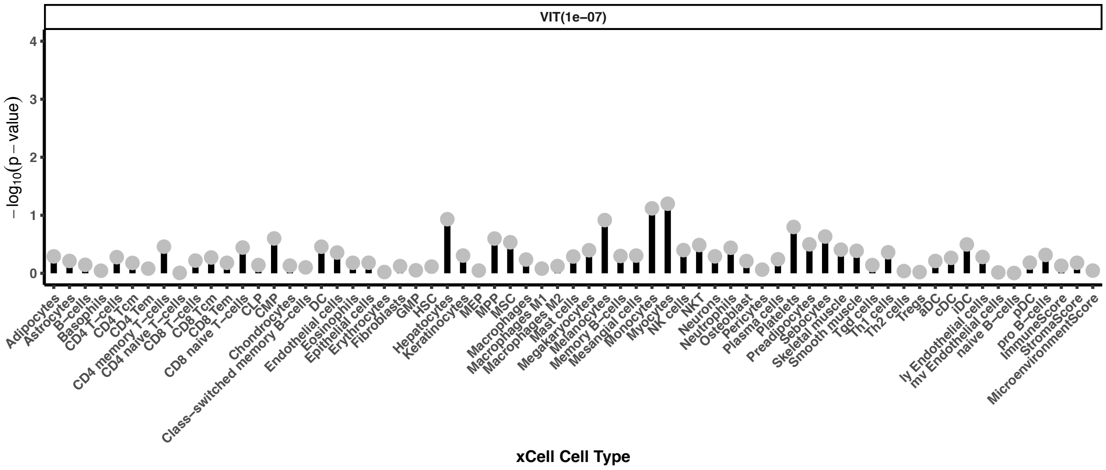


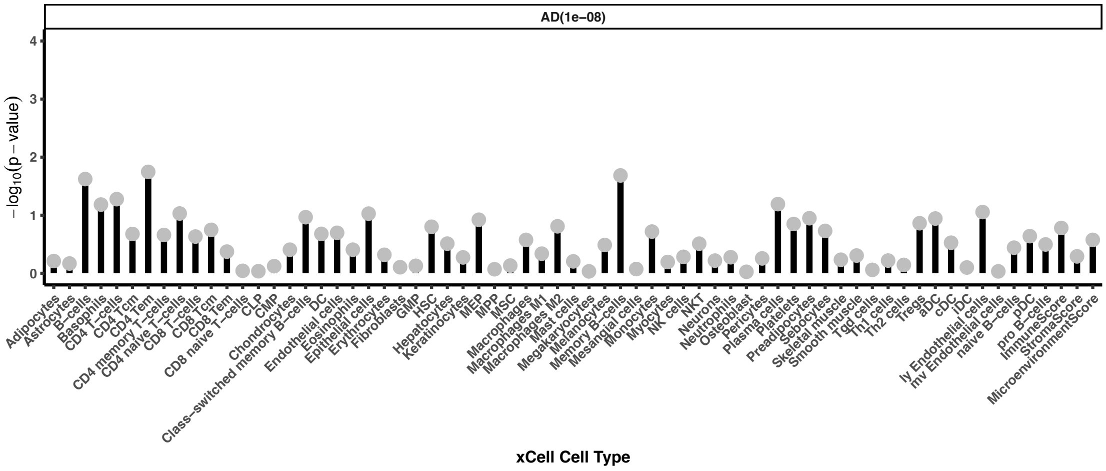


**Figure S10.** Negative $\log_{10}$ p-values testing for an association between xCell cell type enrichment scores and PRSs by linear regression. Tests were adjusted for genotype eigenvectors and gender. xCell enrichment scores were normalized to quantiles of a standard normal distribution. None of the associations were significant at FDR of 10% estimated using the BH procedure accounting for the total number of xCell cell types tested for each PRS.

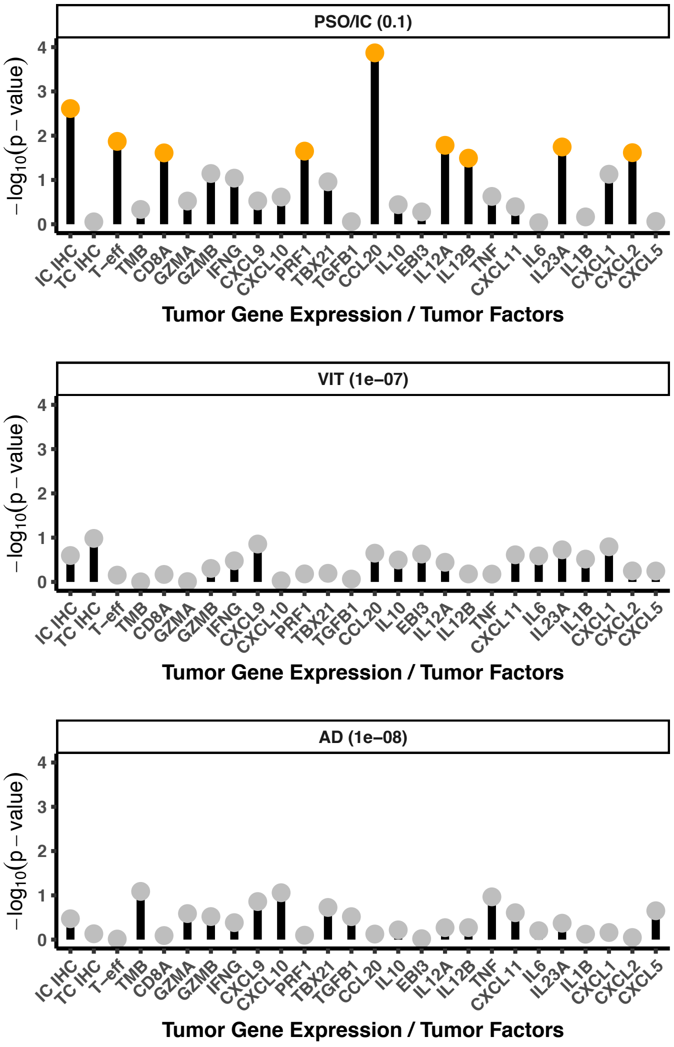


**Figure S11.** Assessing relevance of PRSs in differing tumor immune contexts. Negative $\log_{10}$ p-values for a non-zero 3-way interaction term between PRS (high/low) by trial arm (atezolizumab/chemotherapy) by tumor factor/RNA-seq (high/low) in a Cox proportional hazards model for overall survival. The model controlled for several baseline clinical factors (see **Methods**). High or low was defined on median split of the PRS and median split of tumor factor value. High immune cell and tumor cell IHC for PD-L1 was defined as the IC1/IC2/IC3 and TC1/TC2/TC3 groups respectively. Orange circles designate significant associations at an FDR of 10% estimated using the BH procedure.

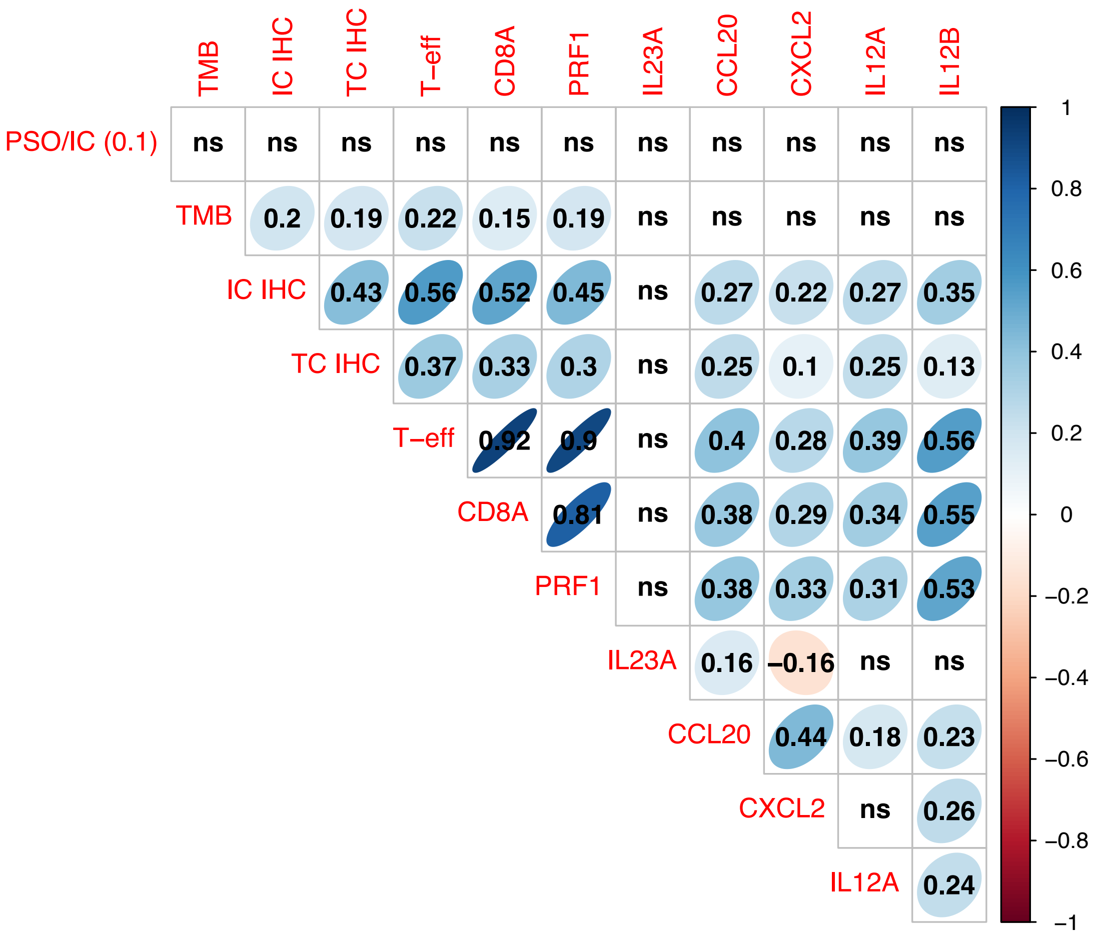


**Figure S12.** Spearman's rank correlation across individuals between the PSO/IC (0.1) PRS and tumor gene expression interactions we identified significant at an FDR of 10%. GWAS p-value cutoff is provided in parentheses. Rank correlations with $p \geq 0.05$ are labeled ns.

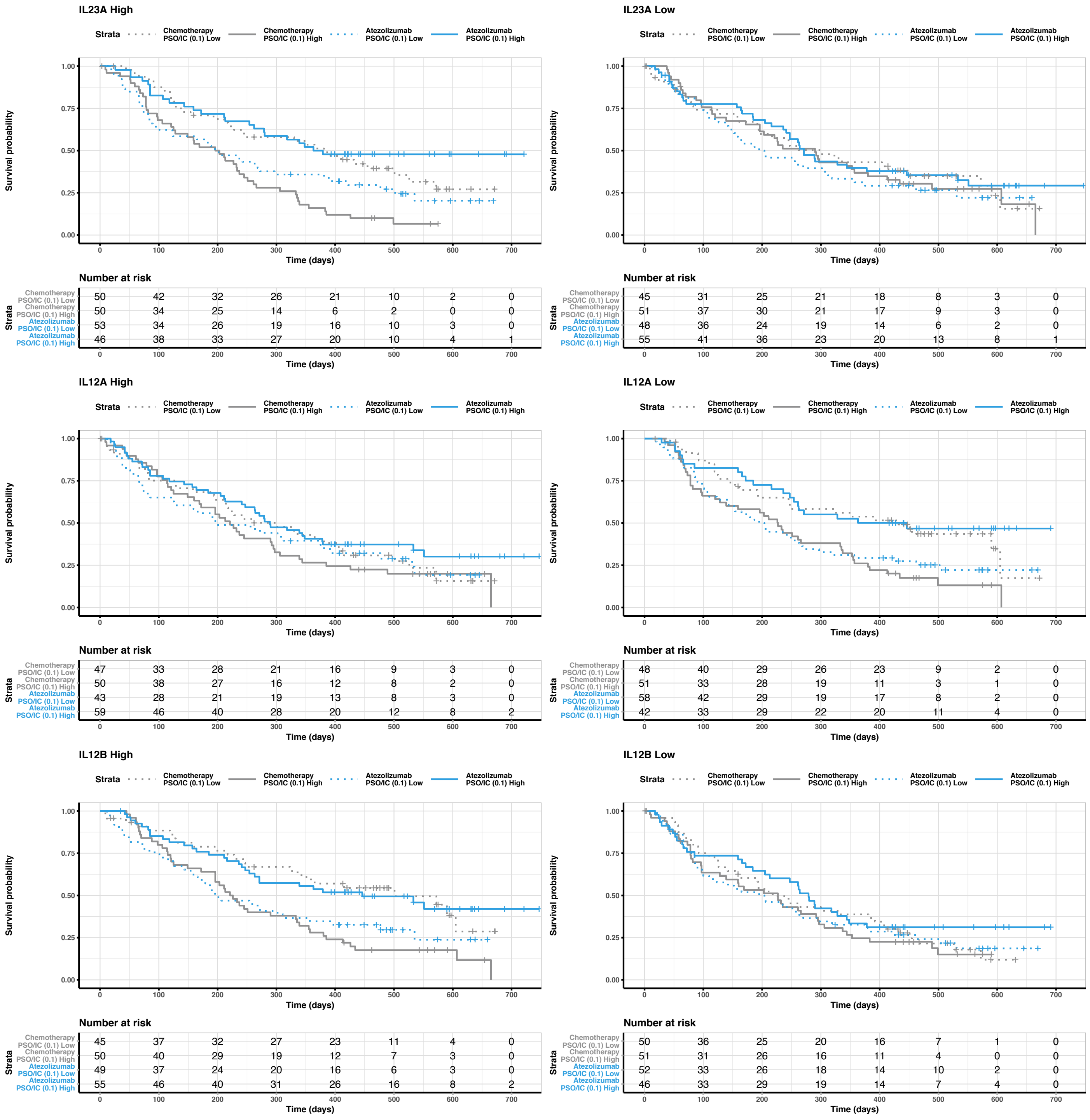


**Figure S13.** Kaplan-Meir plots of overall survival (OS) comparing atezolizumab to chemotherapy for individuals that had high or low genetic risk for psoriasis split also on median tumor gene expression (above median high, left column, and below median low, right column) of IL23A(p19), IL12A(p35), and IL12B(p40). We used the PRS derived from the PSO/IC study at a GWAS p-value cutoff of 0.1 as indicated by the legend above the KM plot. Atezolizumab OS data is plotted in blue. Dotted lines show the low psoriasis risk group. Tick marks designate censoring.

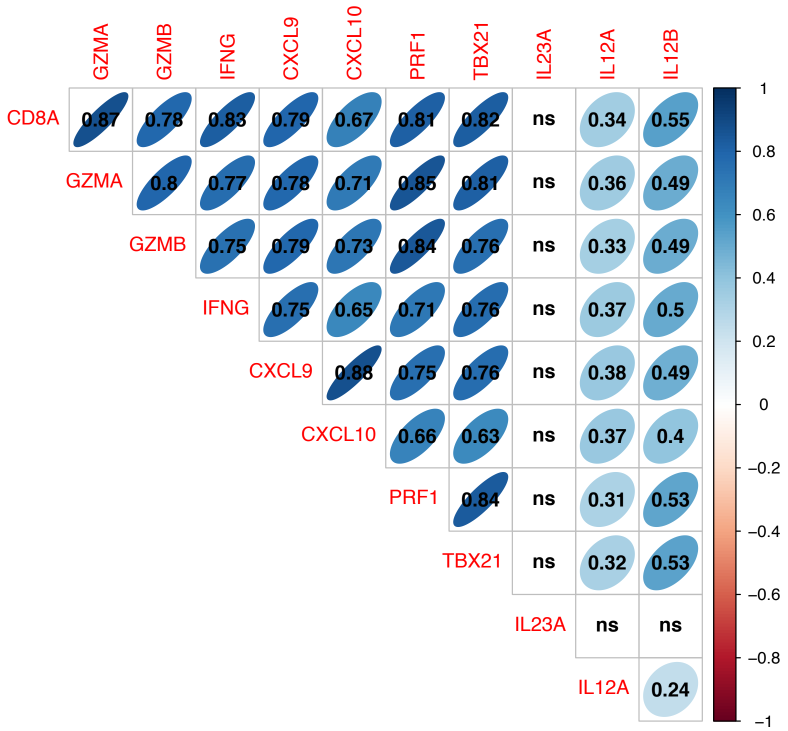


**Figure S14.** Spearman's rank correlation between IL23A, IL12A, and IL12B and CD8$^{+}$ T-effector signature genes across individuals in IMvigor211. Rank correlations with $p \geq 0.05$ are labeled ns.
